# CCDC15 localizes to the centriole inner scaffold and regulates centriole integrity and ciliogenesis

**DOI:** 10.1101/2023.02.16.528810

**Authors:** Melis D. Arslanhan, Emmanuelle Steib, Virginie Hamel, Paul Guichard, Elif Nur Firat-Karalar

## Abstract

Centrioles are evolutionarily conserved microtubule-based organelles critical to form centrosomes and cilia, which act as microtubule-organizing, signaling and motility centers. Biogenesis and maintenance of centrioles with proper number, size and architecture are crucial for their functions during development and physiology. Consequently, their deregulation causes developmental disorders and cancer. Although centriole number control has been extensively studied, less is known about how centrioles are maintained as stable structures with conserved size and architecture over successive cell divisions and upon ciliary and flagellar motility. Here, we addressed this question by identifying and characterizing new components of the centriole inner scaffold, a recently discovered centriolar sub-compartment critical for centriole size control and integrity. To this end, we generated proximity interactomes of Centrin-2 and POC5 and used them to define CCDC15 as a new centriolar protein that co-localizes and interacts with known inner scaffold proteins. Ultrastructure expansion microscopy analysis of CCDC15-depleted cells revealed its functions in centriole length control and integrity, resulting in defective ciliogenesis and Hedgehog signaling. Loss-of-function experiments also defined CCDC15 as a dual regulator for the recruitment of the inner scaffold protein POC1B and the distal SFI1/Centrin complex to the centrioles. Together, our findings uncovered new players and mechanisms of centriole architectural integrity and thereby, provide insights into diseases linked to centriolar defects.

## Introduction

Centrioles are microtubule-based organelles that maintain a conserved number and structure across many eukaryotic cells (Arquint et al., 2014; Azimzadeh and Marshall, 2010; Breslow and Holland, 2019; Brito et al., 2012). They recruit pericentriolar material (PCM) to assemble centrosomes, which act as microtubule-organizing centers in diverse cellular processes including mitotic spindle assembly, cell cycle progression, cell polarity and migration (Bowler et al., 2019; Chavali et al., 2014; Conduit et al., 2015). In quiescent cells, they act as basal bodies to template the formation of the primary cilium, motile cilia or flagella. While the primary cilium is non-motile and functions as a signaling nexus for developmentally important signaling pathways, motile cilia and flagella are required for movement of liquid across specialized epithelia and cell motility (Boutin and Kodjabachian, 2019; Mirvis et al., 2018; Sanchez and Dynlacht, 2016; Wheway et al., 2018). Proper functioning of centrioles during these processes requires spatiotemporal control of their number, size and architecture (Azimzadeh and Marshall, 2010; Loncarek and Bettencourt-Dias, 2018; Nigg and Raff, 2009). Therefore, biogenesis of centrioles is a highly regulated, multistep process and its deregulation is implicated in many human pathologies including cancer, primary microcephaly and ciliopathies (Bettencourt-Dias et al., 2011; Braun and Hildebrandt, 2017; Kathem et al., 2014; Nigg and Raff, 2009; Wang and Dynlacht, 2018; Wheway et al., 2014).

In animal cells, centrioles are composed of nine microtubule triplets radially arranged in a cylinder of about ∼250 nm in diameter and ∼450 nm in length (Chretien et al., 1997; LeGuennec et al., 2021; Loncarek and Bettencourt-Dias, 2018). Although centriole size varies across different species and different cell types in the same species, it is kept relatively constant in cells (Loncarek and Bettencourt-Dias, 2018). Due to their microtubule-based nature, centrioles are inherently polar with microtubule minus ends at their proximal ends and microtubule plus ends at their distal ends (Loncarek and Bettencourt-Dias, 2018). The proximal ends of centrioles contain the cartwheel structure that scaffolds centriole assembly, participates in dictating the diameter of the centriole barrel and imparts its 9-fold symmetry. On the other hand, their distal ends contain appendages that are required for cilium assembly and microtubule anchorage (Guichard et al., 2018; Nakazawa et al., 2007). In contrast to the dynamic cytoplasmic microtubules, centriolar microtubules are exceptionally stable as they resist microtubule depolymerization induced by drug and cold treatment and mitotic entry and exhibit slow turn-over in pulse-chase experiments (Belmont et al., 1990; Bornens et al., 1987; Kochanski and Borisy, 1990; Mitchison and Kirschner, 1986). Importantly, the stable nature of centrioles enables them to withstand mechanical forces during centriole duplication and cell division as well as upon ciliary and flagellar motility.

In most cells, centrioles duplicate precisely only once in early S phase, which involves the formation of a procentriole adjacent to each pre-existing centriole (Carvalho-Santos et al., 2010; Firat-Karalar and Stearns, 2014; Holland et al., 2010). Procentrioles subsequently elongates until mitosis and are then segregated to the daughter cells by the mitotic spindle. Despite our extensive understanding of assembly mechanisms for centrioles, relatively less is known about how their size and architectural integrity is established and maintained. Modifications of tubulin subunits such as acetylation, detyrosination and polyglutamylation was shown to contribute to the stability of the centriolar microtubules (Janke, 2014; Wloga et al., 2017). Moreover, a group of centriolar microtubule-associated proteins has been described for their roles in assembling full-length centrioles. On one hand, CPAP, CEP120 and SPICE act as activators of centriole length via promoting elongation or stabilization of centriolar microtubules (Comartin et al., 2013; Schmidt et al., 2009; Sharma et al., 2021). On the other hand, CP110-CEP97 complex acts as inhibitors by capping the distal ends of centrioles and restricting microtubule growth or depolymerizing/destabilizing centriolar microtubules (Comartin et al., 2013; Franz et al., 2013; Schmidt et al., 2009; Sharma et al., 2021; Spektor et al., 2007). Moreover, the evolutionarily conserved centriole protein POC1 was shown to localize to centriolar microtubules and function in assembling centrioles with proper length and integrity in human cells, zebrafish, *Drosophila melanogaster* and *Tetrahymena thermophilia* (Blachon et al., 2009; Keller et al., 2009; Pearson et al., 2009; Venoux et al., 2013). Finally, Delta- and Epsilon-tubulin were also described in several organisms for their roles in stable centriole formation and inheritance via maintaining triplet microtubules (Dutcher et al., 2002; Dutcher and Trabuco, 1998; Garreau de Loubresse et al., 2001; O’Toole et al., 2003).

Recent advances in expansion microscopy and cryo-tomography have resulted in discovering the inner scaffold in the central region of the centriole. It has been characterized as a regulator of centriole length and integrity (Atorino et al., 2020; Le Guennec et al., 2020; Mercey et al., 2022; Pearson et al., 2009; Schweizer et al., 2021; Steib et al., 2020). Cryo-tomography analysis of the inner scaffold in *Paramecium teraurelia*, *Chlamydomonas reinhardtii, Naegleria gruberi,* and human centrioles defined it as an evolutionarily conserved structural feature that forms a periodic, helical structure composed of repeating units of scaffold protein complexes (Le Guennec et al., 2020). Ultrastructure Expansion Microscopy (U-ExM) analysis of the centrioles revealed nanoscale organization of POC5, POC1B, FAM161A, Centrin-2, WDR90, gamma-TURC and HAUS6 at the centriole lumen (Hamel et al., 2017; Le Guennec et al., 2020; Schweizer et al., 2021; Steib et al., 2020). Although POC5, POC1B and FAM161A localization was restricted to the central region of the centriole, Centrin-2 was shown to localize both to the distal and central regions. POC5, POC1B, FAM161A and Centrin-2 form a complex that binds to microtubules via FAM161A and another microtubule-associated protein WDR90 connects the inner scaffold to the microtubule triplets of the centrioles (Le Guennec et al., 2020; Steib et al., 2020). Centrin-2 also forms a complex with SFI1 at the distal end (Laporte et al., 2022). Loss-of-function studies defined functions for WDR90, augmin and POC5, and thereby for the inner scaffold, during centriole length regulation and stabilization of the centriole architecture (Schweizer et al., 2021). The distal SFI1/Centrin complex was also described for their roles in conferring stability on centrioles. Disruption of the inner scaffold and the distal SFI1/Centrin complex also resulted in defective primary cilium formation. Consistent with this phenotype, mutations affecting POC1B, POC5, Centrin-2 and FAM161A were linked to inherited retinal degeneration caused by loss of photoreceptor cells (Langmann et al., 2010; Roosing et al., 2014; Weisz Hubshman et al., 2018; Ying et al., 2019). Recent work in photoreceptor cells characterized the inner scaffold within their connecting cilium that lines the inner wall of the microtubule doublets and maintains their cohesion (Mercey et al., 2022).

Cellular functions and disease links of the inner scaffold highlight its functional significance in cells and organisms. However, the full repertoire of inner scaffold proteins involved in this regulation and the mechanisms by which they regulate centriole size is not well understood. Applying the U-ExM method to investigate known centrosome proteins has started to define the composition of the inner scaffold. However, its targeted nature limited the identification of the full repertoire of inner scaffold proteins. Here, we used the known inner scaffold proteins as probes to identify the molecular makeup of the inner scaffold in an unbiased way. To this end, we generated proximity interaction maps for POC5 and Centrin-2 and used these maps to define CCDC15 as a new centriolar protein that co-localizes and interacts with known inner scaffold proteins. Depletion of CCDC15 resulted in defective recruitment of the inner scaffold protein POC1B and distal end SFI1/Centrin complex. Consequently, CCDC66 loss compromised centriole size and integrity, wherein the basal body had a reduced ability to form primary cilium. The cilia that formed in CCDC15-depleted cells were shorter and defective in responding to Hedgehog stimuli. Our findings identify CCDC15 as a new regulator of centriole size and architectural integrity and thereby, for maintaining the ability of centrioles to template the assembly of functional cilia.

## Results

### Proximity mapping at the centriole inner scaffold identifies new centriolar proteins

To characterize new components of the centriole inner scaffold, we identified the proximity interactome of Centrin-2 and POC5 in human embryonic kidney (HEK293T) cells using the BioID approach. To this end, we generated cells that stably express V5BirA* (hereafter BirA*) fusions of POC5 and Centrin-2 and validated them by blotting and staining for V5 to detect the fusion protein, streptavidin to detect biotinylated proteins, and/or gamma-tubulin to mark the centrosome (Fig. 1A, 1B). BirA*-fusions of Centrin-2 and POC5 both localized to the centrosome, and stimulated biotinylation at the centrosome (Fig. 1A). Immunoblotting cell lysates confirmed the expression of the fusion proteins and streptavidin pulldown of the biotinylated proteins including the bait proteins POC5 and Centrin-2 (Fig. 1B).

**Fig. 1.**
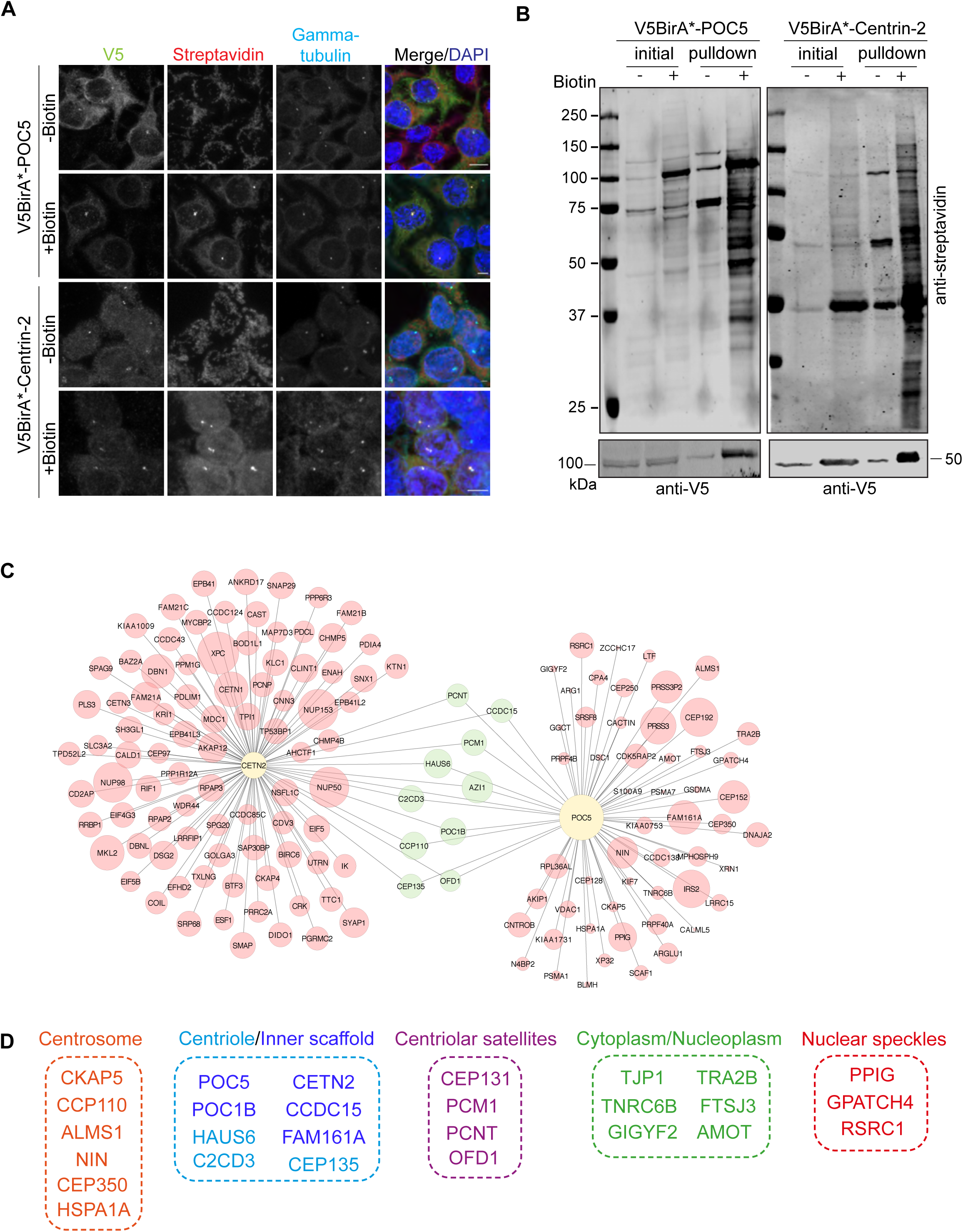
Identification of POC5 and Centrin-2 proximity interactome. **A.** Biotinylation of the centrosome by V5BirA*-POC5 and V5BirA*-Centrin-2. HEK293T cells stably expressing V5BirA*-POC5 and V5BirA*-Centrin-2 were treated with biotin for 18 h. Cells were then fixed and stained for the protein of interest with V5, biotinylated proteins with fluorescent streptavidin and centrosome with anti-gamma-tubulin antibody. DNA was visualized with DAPI. Scale bar, 10 μm **B.** HEK293T cells stably expressing V5BirA*-POC5 and V5BirA*-Centrin2 were lysed, and biotinylated proteins were precipitated by streptavidin beads. The initial sample (initial) and immunoprecipitated biotinylated proteins (pulldown) were run on a gel and immunoblotted with fluorescent coupled streptavidin and V5 antibodies. **C.** POC5 and Centrin-2 proximity interactome map. High confidence proximity interactors of POC5 and Centrin-2 were determined by using NSAF analysis. The interactome map containing the first 100 proximity interactome of Centrin-2 and all the proximity interactors POC5 was drawn in CytoScape and the shared interactome were visualized in green circles. The circle size corresponds to the fold change. **D.** Cellular compartment analysis of the shared proximity interactors of Centrin-2 and POC5. The GO analysis of the shared proximity interactome was determined using DAVID.

For large-scale pulldowns, asynchronous stable cells were grown in 5×15 cm plates and incubated with 50 μM biotin for 18 h. Following denaturing lysis, biotinylated proteins were precipitated by streptavidin beads and analyzed by mass spectrometry. Four biological replicates for POC5 and three biological replicates for Centrin-2 fusions were analyzed by mass spectrometry. As controls, we used mass spectrometry data from four biological replicates derived from control V5BirA* cells (Arslanhan et al., 2021). High-confidence proximity interactomes of POC5 and Centrin-2 were defined by filtering out low-confidence interactors using three different analysis methods and thresholds as described in the methods (Table S1). First, we performed Normalized Spectral Abundance Factor (NSAF) analysis and included proteins with log^2^ NSAF value greater than 1 (Firat-Karalar et al., 2014; Zybailov et al., 2006). Second, we accounted for proteins identified in at least 2 replicates. Third, we removed common mass spectrometry contaminants (>30% of the contaminants) by using The Contaminant Repository for Affinity Purification – Mass Spectometry data (CRAPome) (Mellacheruvu et al., 2013). Altogether, these thresholds yielded 68 and 480 proteins as high confidence interactors of POC5 and Centrin-2, respectively (Table S1, Fig. S1)

To validate the proximity interactomes of Centrin-2 and POC5, we performed Gene Ontology (GO) enrichment analysis based on their “biological process” and “cellular compartment” (Table S2). As shown in Fig. S1, Centrin-2 and POC5 proximity interactomes were enriched for GO categories that are relevant for their published functions during centrosomal and/or nuclear biological processes and related cellular compartments (Azimzadeh et al., 2009; Dantas et al., 2013; Heydeck et al., 2020; Khouj et al., 2019; Resendes et al., 2008; Salisbury et al., 2002; Steib et al., 2020; Yang et al., 2010; Ying et al., 2019). For example, the highly enriched GO categories for biological processes include cell division, protein folding, mRNA splicing, and chromatin remodeling for Centrin-2 and centriole replication, protein localization to centrosome and cilium assembly for POC5 (Fig. S1A-D). Moreover, GO term analysis of cellular compartment showed significant enrichment for centriole, centrosome, nuclear speckles and centriolar satellites for both Centrin-2 and POC5 (Fig. S1A-D). In addition to GO analysis, we ranked POC5 and Centrin-2 proximity interactors by their fold change into an interaction network, combining STRING database and ClusterONE plug-in on Cytoscape (Fig. S1E, S1F). The resulting network identified a diverse array of proteins from five major functional clusters (*P* < 0.005) for Centrin-2 and two major clusters for POC5. These clusters include nucleoplasm, cell junctions, mitotic spindle and spindle pole, nucleolus and centriole and basal body for Centrin-2, and centrosome and extracellular exosome for POC5 (Fig. S1E, S1F).

To identify new centriole proteins, we focused on the proximity interactors shared by POC5 and Centrin-2 datasets, which included 27 proteins from five major cellular compartments from the centriole/inner scaffold, centrosome, centriolar satellites, cytoplasm/nucleoplasm and nuclear speckles (Fig. 1C, D). The identification of previously characterized centriole proteins FAM161A, POC1B, HAUS6, CEP135 and C2CD3 validated our interactomes as resources for future studies. Among the shared interactors, we chose CCDC15 for further cellular characterization as it had been identified in the centriolar satellite proteome and in proximity maps of centrosome/centriolar satellite proteins including the ones involved in biogenesis of centrioles (i.e. PLK4, KIAA0753) (Firat-Karalar et al., 2014; Gheiratmand et al., 2019; Gupta et al., 2015; Quarantotti et al., 2019). CCDC15 was also shown to be implicated in tumorigenesis (Tang et al., 2020).

### CCDC15 localizes to centrioles throughout the cell cycle

We investigated the localization of endogenous CCDC15 and mNeonGreen (mNG)-CCDC15 fusion protein in human Retinal Pigment Epithelium 1 (RPE1) cells (Fig. 2, S2). Antibodies against CCDC15 and the centriole marker Centrin-2 showed that CCDC15 localizes to the centrosome throughout the cell cycle (Fig. 2A). CCDC15 was detected as two foci per cell in interphase and four foci per cell from G2 through mitosis, showing that it is a centriole protein (Fig. 2A). Of note, centrosomal levels of CCDC15 remained unaltered upon nocodazole-induced microtubule depolymerization (Fig S2A, S2B). Analogous to its endogenous localization, transiently expressed mNG-CCDC15 localized to the centrosomes (Fig. S2C). In a subset of transfected cells, mNG-CCDC15 localized to centriolar satellites marked by PCM1 (Fig. S2D). Given the role of centriolar satellites in centrosomal protein targeting, we quantified centrosomal levels of CCDC15 in RPE1::PCM1^-/-^ satellite-less cells and found that it was reduced relative to control cells (18.5% reduction, p<0.0001) (Fig. S2E, 2F) (Odabasi et al., 2020). These results suggest that centriolar satellites contribute to maintaining centrosomal CCDC15 levels.

**Fig. 2.**
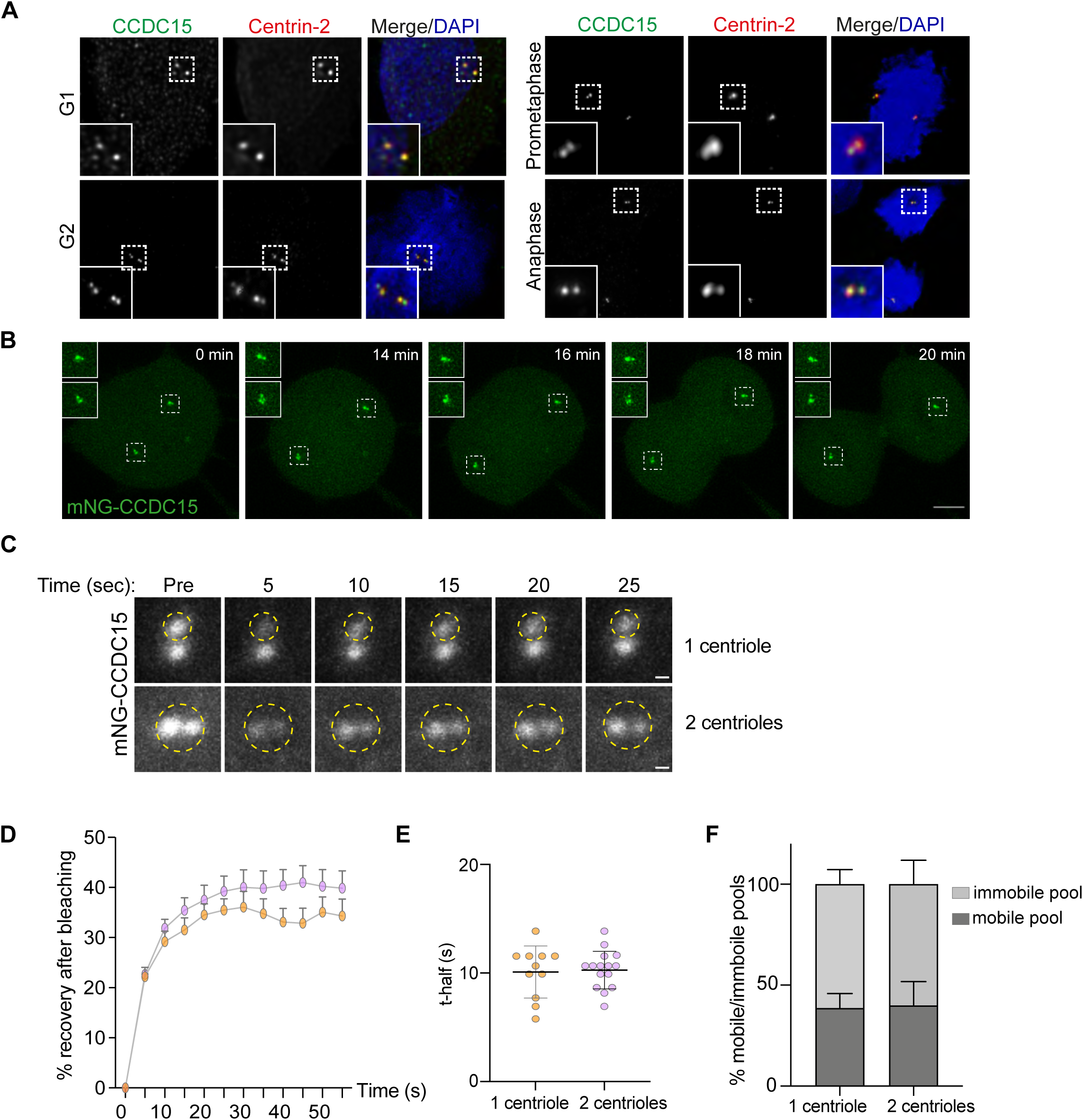
CCDC15 is stably associated with centrosomes throughout the cell cycle. **A.** Localization of endogenous CCDC15 to the centrioles in different cell cycle stages. RPE1 cells were stained with antibodies against the centriole marker Centrin-2 and CCDC15. DNA was visualized with DAPI. Scale bar, 5 μm **B.** Spatiotemporal dynamics of CCDC15 during cell cycle. RPE1 cells stably expressing mNeonGreen-CCDC15 during mNeonGreen-CCDC15 were imaged every 2 minutes. Scale bar, 1 μm **C.** Fluorescence recovery after photobleaching (FRAP) analysis of CCDC15 dynamics at centrioles. RPE1 cells stably expressing mNeonGreen-CCDC15 were grown in a glass-bottom dish, centrioles indicated with yellow circle (3 μm^2^) were photobleached and then were assessed at the indicated times after photobleaching. **D.** Percentage of recovery graph of (C). Individual FRAP experiments from two biological replicates were fitted into a one phase association curves. n=11 for one centriole curve and n=15 for two centrioles curve. **E.** Half-time analysis of were calculated using recovery data from (D). Error bars, SD **F.** Percentage of mobile and immobile pools of CCDC15 at centrioles were calculated from (D). Error bars, SD

We examined CCDC15 dynamics during cell division by time-lapse imaging of RPE1 cells that stably express mNG-CCDC15 (Fig. 2, Movie S1). Consistent with its centriolar localization, mNG-CCDC15 localized to four foci in dividing cells. To investigate the dynamics of CCDC15 association with the centrioles, we photobleached one or both centrioles of in interphase cells and quantified fluorescence recovery over time (Fig. 2D, S2G, S2H). In both cases, only ∼40% of the mNG-CCDC15 fluorescence recovered rapidly (halftime of ∼10 sec), identifying the remaining 60% as the stably associated CCDC15 pool (Fig.2E, 2F). This result shows that the majority of CCDC15 is stably associated pools with the centrosome and is analogous to the behavior of other centriolar proteins such as RTTN and POC1B (Sydor et al., 2018; Venoux et al., 2013).

To determine the sub-centrosomal localization of CCDC15, we examined its localization relative to the markers of the centriole distal end lumen (Centrin-3), distal appendages (CEP164), centriole proximal end linker (Rootletin), centriole microtubule wall (polyglutamylated tubulin) and PCM (gamma-tubulin). CCDC15 localized to the central region of centrioles between the distal marker CEP164 and the proximal end markers Rootletin and gamma-tubulin (Fig. 3A). Given the proximity interactions of CCDC15 with inner scaffold proteins, we next studied their potential co-localization and interaction. To this end, we analysed CCDC15 localization in cells transfected with GFP-FAM161A, GFP-POC1B and V5-POC5 (Fig. 3B). In all cases, CCDC15 partially co-localized with these proteins at the centriole. We also performed co-immunoprecipitation experiments in cells co-expressing GFP-CCDC15 or FLAG-CCDC15 with V5-POC5, GFP-POC1B, FLAG-FAM161A and GFP-Centrin-2. CCDC15 co-precipitated with POC5, Centrin-2, POC1B and FAM161A, but not with the negative controls (Fig. 3C-F). CCDC15 neither interacted with SASS6, a centriole protein that does not localize to the inner scaffold (Fig. 3G). Taken together, our findings from localization and interaction experiments define CCDC15 as a new centriole core protein.

**Fig. 3.**
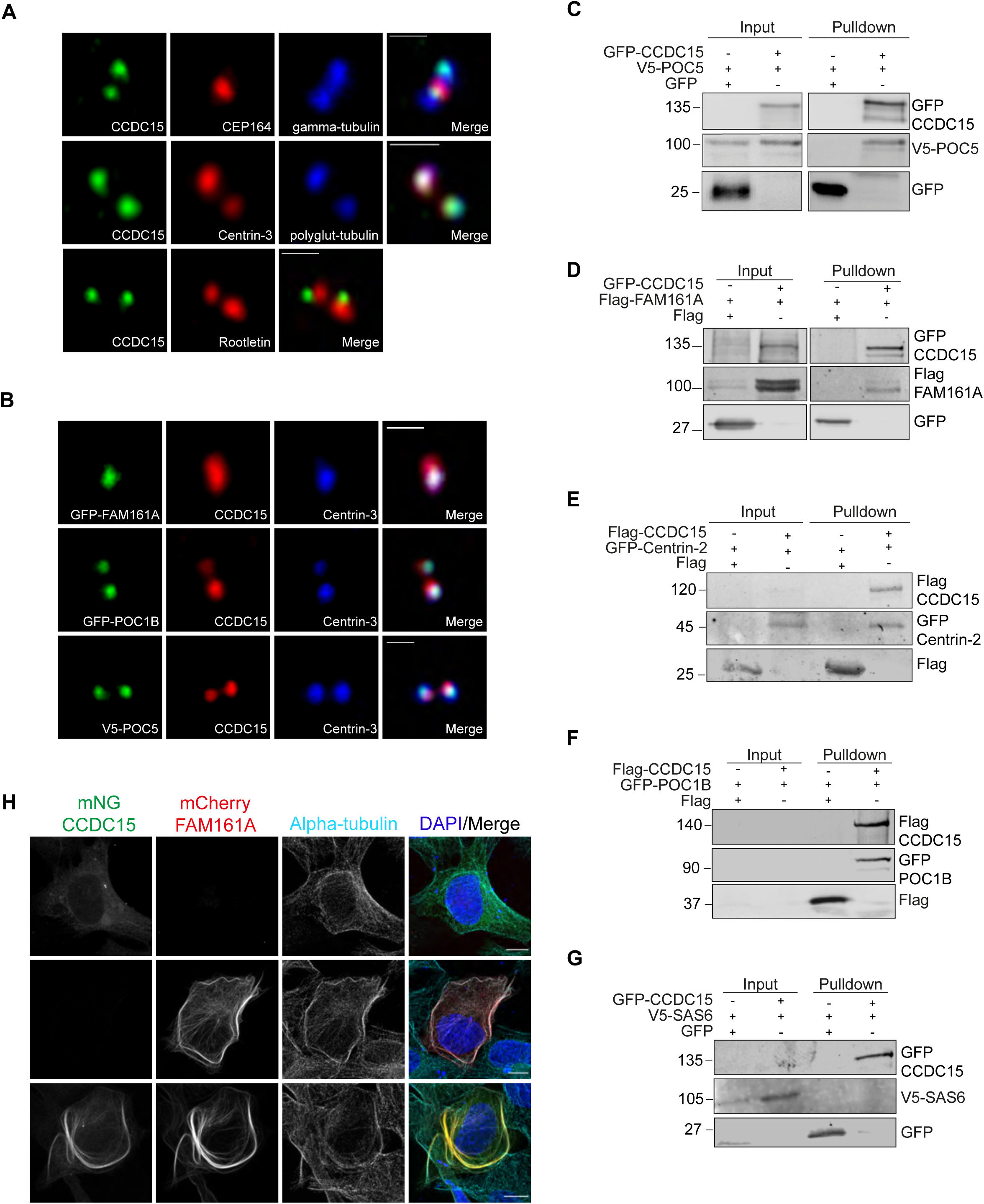
CCDC15 localizes to the centrioles and interacts with inner scaffold proteins. **A.** Localization of CCDC15 relative to proximal and distal end markers of centrioles. RPE1 cells were fixed with methanol and stained for CCDC15 and markers for distal appendages (CEP164), proximal end linker (Rootletin), centriole microtubule wall (polyglutamylated tubulin), centriole distal end lumen (Centrin-3) and PCM (gamma-tubulin). Scale bar, 1 μm **B.** Co-localization analysis of CCDC15 with known inner scaffold proteins. U2OS cells were transiently transfected with GFP-FAM161A, GFP-POC1B and V5-BirA*-POC5 and stained with antibodies against CCDC15, Centrin-3 and GFP or V5 to mark the inner scaffold proteins. Scale bar, 1 μm **C-F.** CCDC15 interacts with (C) POC5, (D) FAM161A, (E) Centrin-2 and (F) POC1B. HEK293T cells were transfected with the indicated plasmids. 24 h after transfection, cell lysates were collected and CCDC15 was precipitated using GFP-binding protein (GBP) beads. The initial sample and immunoprecipitated proteins were run on a gel and immunoblotted with GFP, Flag and/or V5 antibodies. **G.** CCDC15 does not interact with the proximal end protein SAS6. HEK293T cells were transfected with GFP-CCDC15 and V5BirA*-SAS6. 24 h after transfection, cell lysates were collected and CCDC15 was precipitated using GBP beads. The initial sample and immunoprecipitated proteins were run on a gel and immunoblotted with GFP and V5 antibodies. **H.** Displacement of CCDC15 to the microtubules upon co-expression with FAM161A. U2OS cells were transfected with only mNeonGreen-CCDC15, mCherry-FAM161A or both. Cells were fixed with methanol and stained with antibodies against the epitope tags and the microtubule marker alpha tubulin. Scale bar, 10 μm

Given that inner scaffold ensures stability of centrioles via binding to centriolar microtubules, we also investigated the nature of CCDC15 interaction with microtubules. The microtubule-associated protein FAM161A was shown to act as a scaffold to recruit inner scaffold proteins POC5 and POC1B to the microtubules (Le Guennec et al., 2020). Therefore, we hypothesized that CCDC15 depended on FAM161A for its recruitment to microtubules. To test this, mNG-CCDC15 and mCherry-FAM161A were expressed alone or in combination in U2OS cells (Fig. 3H). Although inner scaffold proteins tend to self-associate, mNG-CCDC15 did not induce the formation of filamentous structures and localized to centrioles when expressed alone (Fig. 3H). Strikingly, its co-expression with FAM161A resulted in its redistribution to microtubules (Fig. 3H). This result suggests that CCDC15 might form a microtubule-associated complex with inner scaffold proteins.

### CCDC15 is a component of the centriole inner scaffold

For nanoscale mapping of CCDC15 localization inside the centriole, we analyzed its distribution using Ultrastructural Expansion Microscopy (U-ExM) (Gambarotto et al., 2019). In RPE1 cells immunostained with antibodies against CCDC15 and tubulin, we found that CCDC15 localizes to the inner core of the centrioles shown in longitudinal and top views (Fig. 4A, 4B). From longitudinal views, we calculated centrioles length as represented by tubulin staining as 446.7 nm (± 45.29 nm) and CCDC15 length inside the centriole as 250.1 (± 41.11 nm) (Fig. 4C). This result indicates that CCDC15 spans 55%± 4.08 nm of the centrioles (Fig. 4D). From the top views, we calculated the average distance between CCDC15 and tubulin maximum intensity signal from the exterior to the interior of the centriole and found that it was *Δ* = 17 ± 2 nm, showing that CCDC15 signal is shifted toward the centriole lumen relative to the tubulin signal (Fig. 4E). This is consistent with the localization profile of other inner scaffold proteins. Furthermore, the comparison of their relative shifts suggests that CCDC15 might localize between POC1B and FAM161A (Le Guennec et al., 2020).

**Fig. 4.**
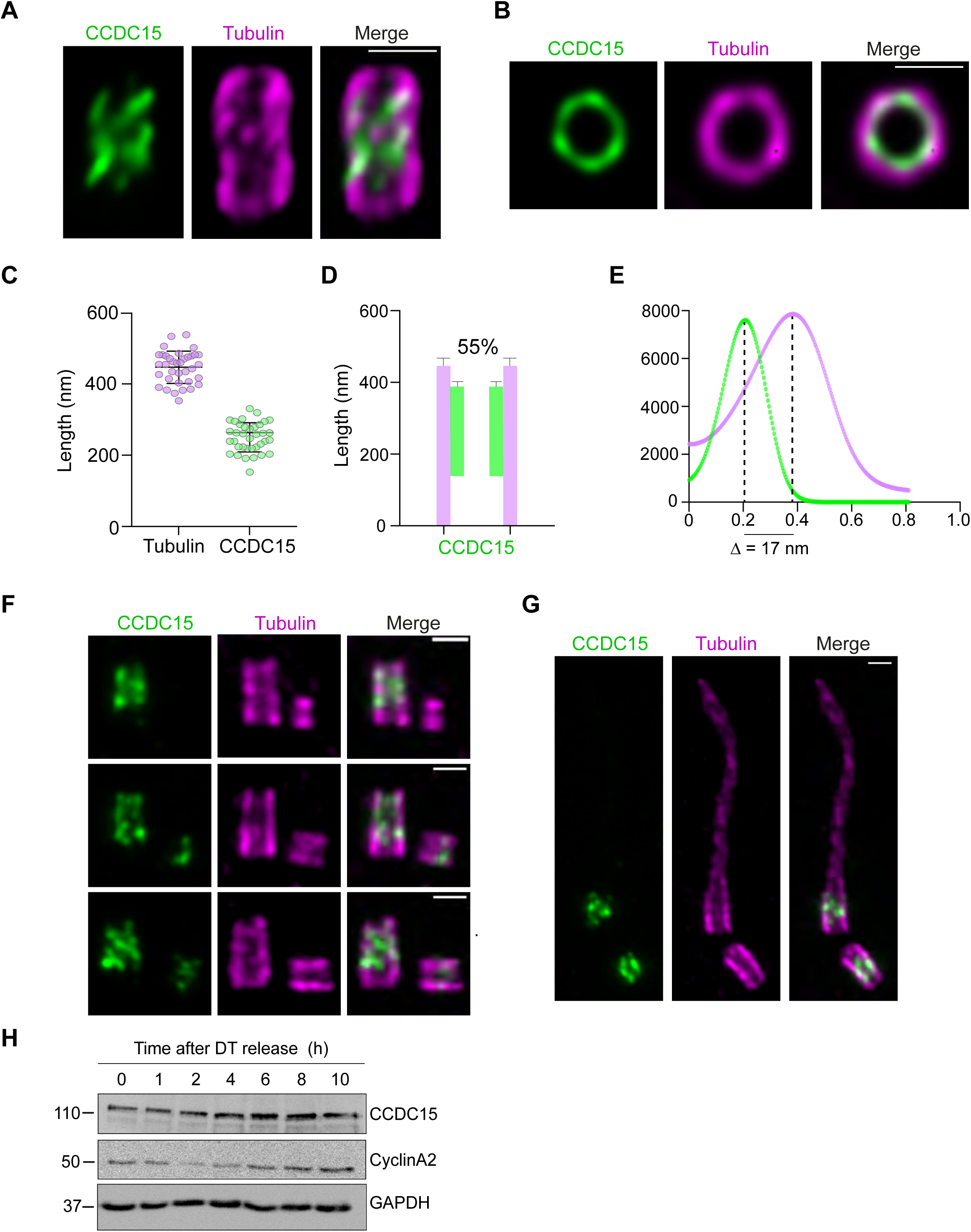
CCDC15 localizes to the inner scaffold. **A.** Representative confocal images of expanded RPE1 mature centrioles expanded using U-ExM and stained for tubulin (magenta) and CCDC15 (green). Scale bar, 1 μm **B.** Top views confocal images of RPE1 mature centriole images in U-ExM stained for tubulin in magenta and CCDC15 in green. Scale bar, 1 μm **C.** Respective lengths of tubulin and CCDC15 based on (A). Error bars, SD: Tubulin, 446 nm ± 45, CCDC15, 250 nm ± 41 nm; n>15 cells from two independent experiments. **D.** Position of CCDC15 along the centriole with its respective percentage of centriole coverage, which was calculated as 55% based on (C). **E.** Plot profile of CCDC15 (green) and tubulin (magenta). The distance between the tubulin and CCDC15 rings were calculated as 17 nm ± 2. **F.** Timing of CCDC15 centriolar recruitment during centriole duplication. Representative confocal images of RPE1 centrioles expanded using U-ExM and stained for tubulin (magenta) and CCDC15 (green). Scale bar, 1 μm **G.** Representative confocal images of CCDC15 localization (green) at the basal bodies (magenta) in RPE1 cells serum starved for 24 h and expanded using U-ExM. Scale bar, 1 μm **H.** Expression profile of CCDC15 in synchronized cells. Lysates were run on western blot and immunoblotted with CCDC15, CyclinA2 and GAPDH antibodies. U2OS cells were synchronized at the G1/S transition using a double thymidine (DT) block, then released into the cell cycle. Lysates prepared from cells at different time points were immunoblotted for CCDC15, Cyclin A2 (marker for the G2/M phase) and GAPDH (loading control)

During centriole duplication, a procentriole grows orthogonally to the proximal end of a pre-existing centriole during early S phase, prior to reaching its full length by the following cell cycle (Kong et al., 2020; Loncarek and Bettencourt-Dias, 2018). Inner scaffold proteins including POC5 and WDR90 are recruited to procentrioles in early G2 and reach full incorporation by the end of G2 (Azimzadeh et al., 2009; Steib et al., 2020). To investigate the timing of CCDC15 centriolar recruitment during centriole biogenesis, we examined CCDC15 localization relative to the length of procentrioles that represent cells at different stages of centriole duplication (Fig. 4F) (Hamel et al., 2017; Kong et al., 2020). While CCDC15 did not localize to centrioles in early stages of centriole duplication, it was recruited to the central core in elongated centrioles (Fig. 4F). In cells that formed primary cilia upon serum starvation, CCDC15 localized to the central core of their basal bodies (Fig. 4G). In parallel to localization experiments, we quantified the protein expression profile of CCDC15 in cells synchronized at the G1/S transition using a double thymidine block, then released into the cell cycle. Lysates prepared from cells at different time points were immunoblotted for CCDC15 and Cyclin A2, a cyclin marker for the G2/M phase (Ding et al., 2018; Silva Cascales et al., 2021). CCDC15 levels increased gradually form early S phase until mitosis (0-8 h) (Fig. 4H). These results demonstrate that CCDC15 is a cell cycle-regulated protein recruited during procentriole assembly and that timing of its centriolar incorporation correlates with that of other inner scaffold proteins.

### CCDC15 regulates centriole length and structure

To investigate CCDC15 functions at centrioles, we performed siRNA-mediated loss-of-function experiments and phenotypically characterized CCDC15-depleted cells by imaging-based assays. As assessed by immunofluorescence, CCDC15 was efficiently depleted from RPE1 cells 96 h after transfection with an siRNA against CCDC15 (Fig. S3A). Centrosomal CCDC15 levels were reduced by about 70% (siControl:1± 0.43, siCCDC15:0.31± 0.29) in CCDC15-depleted cells as compared to control cells (Fig. S3A). We also used U-ExM to determine the extent of CCDC15 loss from centrioles transfected with CCDC15 siRNA. About 49.6% (+- 3.96) of the centrioles still had CCDC15 fluorescence signal at one of the centrioles upon CCDC15 siRNA treatment (Fig. 5A, 5B). The inefficient depletion of the mature centriole pool of CCDC15 is analogous to what was observed upon depletion of other centriole lumen and inner scaffold proteins including WDR90 and HAUS6 (Schweizer et al., 2021; Steib et al., 2020).

**Fig. 5.**
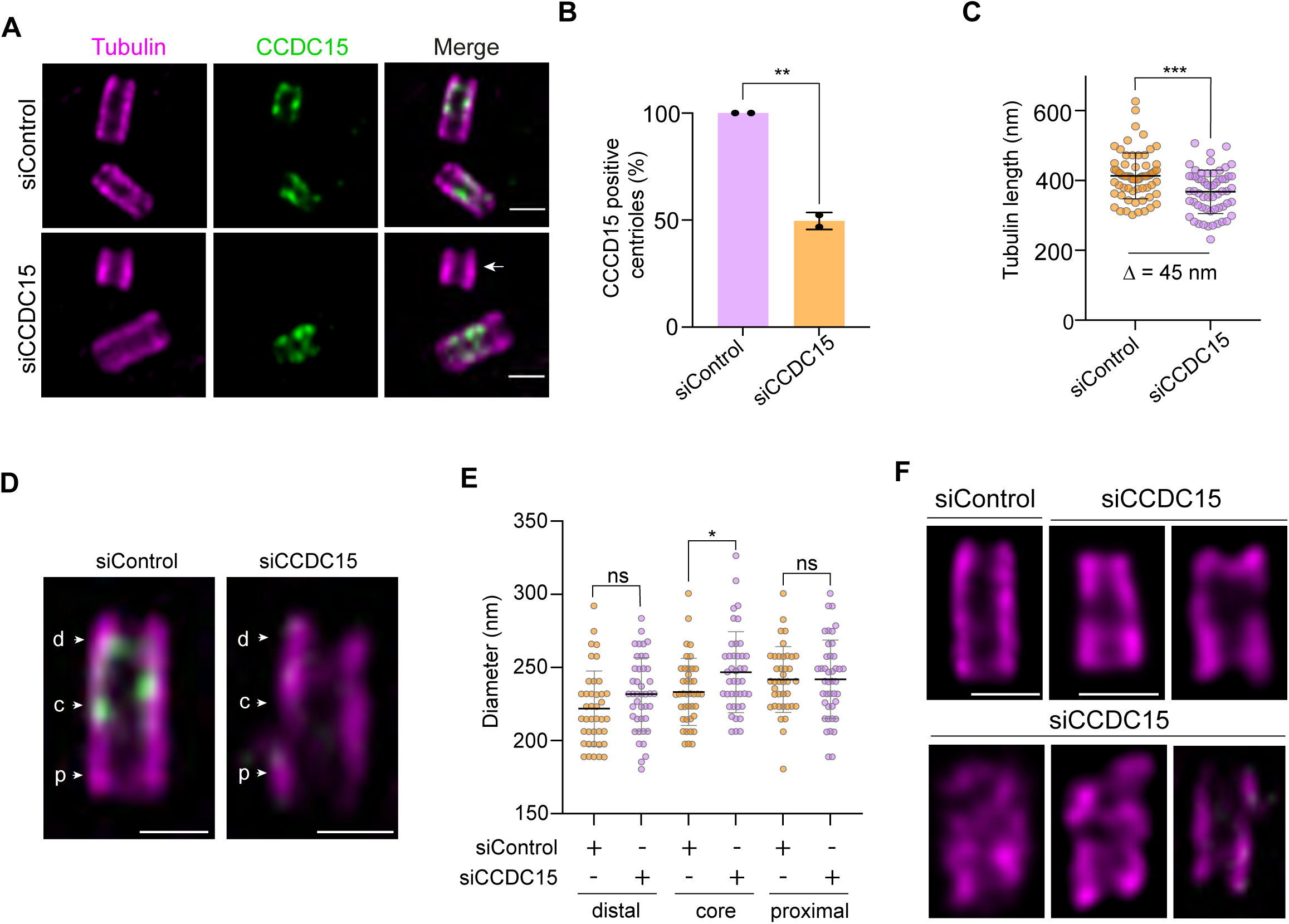
CCDC15 depletion leads to shorter and structurally aberrant centrioles. **A.** CCDC15 depletion leads to shorter centrioles. Representative confocal images of expanded centrioles from control and CCDC15-depleted cells stained for CCDC15 (green) and tubulin (magenta). RPE1 cells were transfected with control or CCDC15 siRNA. 96 posttransfection, cells were expanded by U-ExM. Note that CCDC15 was efficiently depleted from only one of the centrioles in most cells (white arrow). **B.** Quantification of percentage of CCDC15-positive centrioles based on (A). n>40 per experiment. Data represents mean value from two independent experiments. Error bars, SD. siControl=100%, siCCDC15=50% ± 4. p=0.0031. **C.** Centriole length quantification based on (A). Error bars, SD. n>10 per experiment. Data represents mean value from three independent experiments. siControl: 413 nm ± 66, siCCDC15: 367 nm ± 62, p=0.0001. Centrioles depleted of CCDC15 were 45 nm shorter compared to control centrioles in RPE1 cells. **D.** Changes in diameter in distal (d), central (c), and proximal (p) regions of the centrioles. RPE1 cells were transfected with control or CCDC15 siRNA and expanded by U-ExM 96 posttransfection. Gels were stained with tubulin (magenta) and endogenous CCDC15 (green) antibodies. Scale bar, 1 μm **E.** Quantification of (D). Error bars, SD. n>10 per experiment. Data represents mean value from three independent experiments. Distal region: siControl: 222 nm ± 26, siCCDC15: 232 nm ± 25, p=0.0828; core region siControl: 233 nm ± 23, siCCDC15: 247 nm ± 28, p=0.0204; proximal region siControl: 242 nm ± 23, siCCDC15: 242 nm ± 27. **F.** Representative confocal images of expanded centrioles in control and CCDC15-depleted RPE1 cells stained for CCDC15 (green) and tubulin (magenta). Different types of structural defects of CCDC15-depleted cells were represented, which included centrioles with broken, wider or shorter microtubule walls. Scale bar, 1 μm

Inner scaffold proteins were described for their functions during centriole size integrity and architecture. Therefore, we first investigated how CCDC15 loss affects these processes. Using U-ExM, we calculated centriole length using tubulin length as a proxy and found that CCDC15-depleted centrioles exhibited a slight decrease (about 10%) in centriole length relative to control cells (siControl: 413.1 nm± 65.56 nm, siCCDC15: 367.6 nm± 62.06 nm) (Fig. 5C). To quantify morphological changes, we determined the diameter of their proximal, central and distal regions of the centrioles. Despite a slight increase of the diameter in the central core region, the centriole diameter at the proximal and distal regions remained unaltered (Fig. 5D, 5E). Notably, we also found structurally abnormal centrioles in about 12% of CCDC15-depleted cells, displaying open, broken, wider or shorter microtubule walls (Fig. 5F). Together, these results show that CCDC15 is required for centriole length control and integrity.

To determine the functional consequences of centriole abnormalities associated with CCDC15 depletion, we performed assays to assess cell cycle progression and centriole duplication. Flow cytometry analysis of the asynchronous cells showed that control and CCDC15-depleted cells had similar cell cycle profiles (Fig. S3B). We also quantified centriole number by counting the number of centrin-positive foci in asynchronous cultures and found that CCDC15 had no effect on the number of centrioles (Fig. S3C). We further investigated cartwheel assembly by counting the number of SAS-6 foci in cells positive for nuclear proliferating cell nuclear antigen (PCNA), a marker of DNA replication. The percentage of control and CCDC15-depleted cells with two foci were similar, indicating that CCDC15 is not required for centrosomal SAS-6 recruitment (Fig. S3D, E). However, when the mitotic time was extended to 18 h by treatment with the Eg5 inhibitor STLC, we found that 63.45%±3.77 of CCDC15-depleted cells had the expected number of at least four centrin foci as compared to 95.26%±1.47 of control cells (Fig. S3F, 3G). This result suggests centriole destabilization associated with CCDC15 loss, which was also reported for the centriole lumen protein HAUS6 (Schweizer et al., 2021). In addition to canonical duplication, we tested whether CCDC15 is required for centriole amplification. U2OS cells were transfected with control and CCDC15 siRNAs and treated with hydroxyurea for 48 h to induce centriole amplification. We observed that 42.30%±1.27 control and 27.75%±2.19 of CCDC15-depleted cells had more than 4 centrioles (Fig. S3H, I). We also assayed centriole amplification in RPE1::PLK4 inducible line upon control and CCDC15 treatment. There, 83.05%± 9.05 control and 40.78%± 5.82 of CCDC15-depleted cells of CCDC15-depleted cells had more than four centrioles (Fig. S3J, K). Results from centriole duplication and amplification assays show that CCDC15 is required for efficient centriole amplification, but not for canonical centriole duplication.

### CCDC15 and other inner scaffold proteins cooperate for their recruitment to centrioles

CCDC15 might confer centriole stability on centrioles via regulating the inner scaffold. We tested this hypothesis by analyzing the localization of five known inner scaffold components POC5, POC1B, Centrin-2 and FAM161A in RPE1 cells treated with control and CCDC15 siRNAs. First, we quantified the coverage of these proteins along the centriole by U-ExM in cells stained for the inner scaffold proteins and centriole marker (Fig. 6A, 6B). While the centriolar coverage of POC5 slightly increased, the coverage of POC1B decreased. In contrast, CCDC15 depletion did not alter the central core coverage of Centrin-2 and FAM161A. In addition to their sub-centrosomal mapping by U-ExM, we quantified the centrosomal abundance of inner scaffold proteins by drawing a 3.5 um^2^ ring centered around the centrosome marked by gamma-tubulin or Centrin-2. As compared to control cells, the centrosomal abundance of POC5 increased, POC1B decreased and Centrin-2 and FAM161 remained unaltered upon CCDC15 depletion (Fig. S4A-C). To validate the specificity of CCDC15 functions in recruitment of inner scaffold proteins, we also quantified the centrosomal levels of the PCM protein CEP63 and the distal appendage protein CEP164, and found no statistical differences in their levels between control and CCDC15-depleted cells (Fig. S4D, S4E). The trends in the centrosomal abundance changes were similar to the ones quantified by U-ExM (Fig. 6B). These results show that CCDC15 depletion results in defective recruitment of POC1B to the central core of the centrioles.

**Fig. 6.**
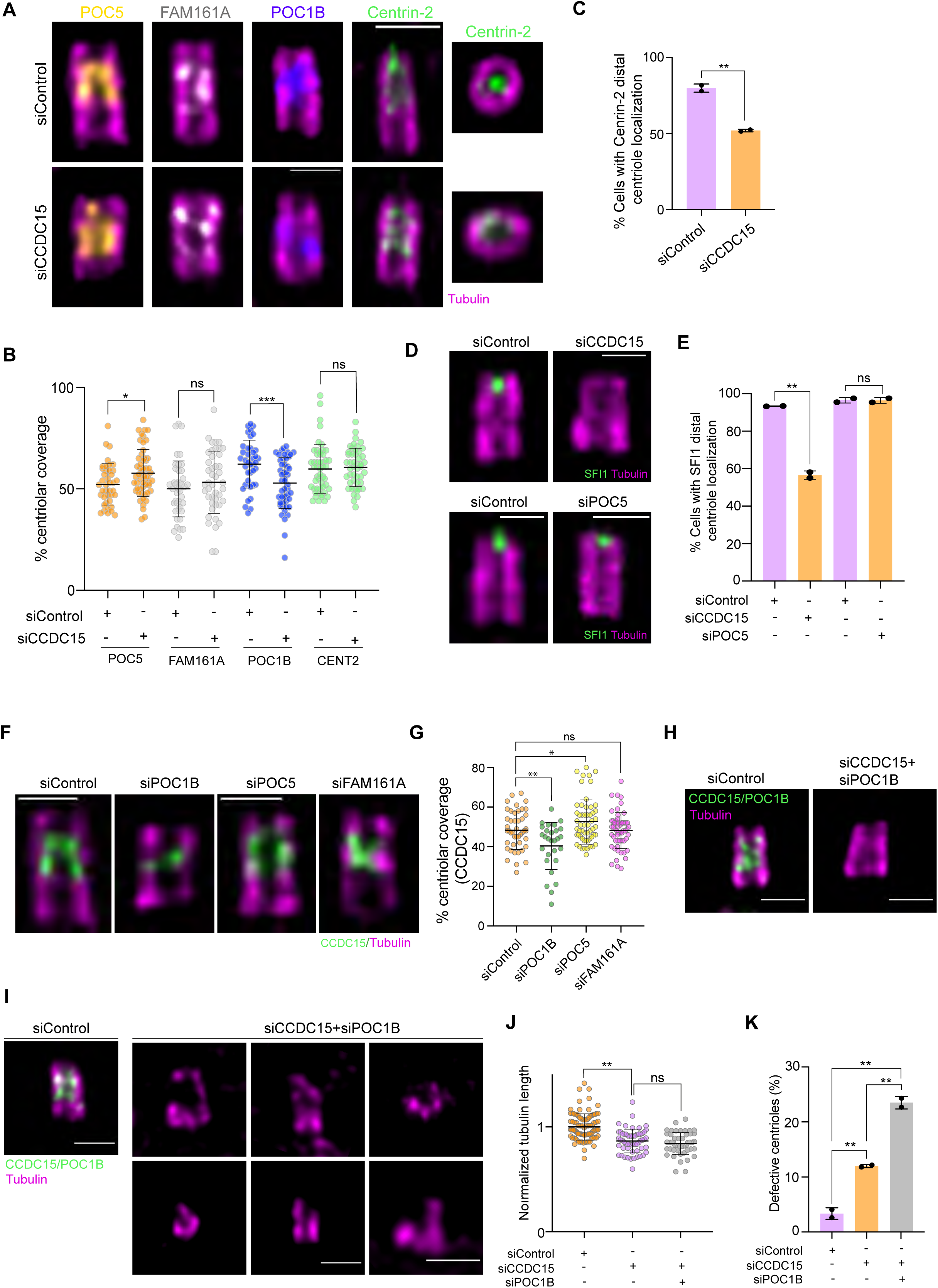
CCDC15 is required for recruitment of inner scaffold proteins to the centriole central core and distal end. **A.** Representative U-ExM images of centriolar core proteins POC5 (orange), FAM161A (grey), POC1B (blue), and Centrin-2 (green) in control or CCDC15-depleted RPE1 cells. Cells were expanded 96 h after siRNA transfection and immunostained with the indicated antibodies for inner core proteins and tubulin (magenta) for centrioles. Top view of centrioles for Centrin-2 was represented in addition to the longitudinal views represented for all proteins. Scale bar, 1 μm **B.** Quantification of coverages of the centriolar proteins in (A). Error bars, SD. n>20 cells per experiment. Data represent mean value from two independent experiments per condition. POC5 coverage: siControl=52% ± 10,siCCDC15=58% ± 12, p=0.0174; FAM161A coverage: siControl=50% ± 14, siCCDC15=53% ± 15, p=0.2981; POC1B coverage: siControl=62% ± 12, siCCDC15=53% ± 12, p=0.0006; Centrin-2 coverage: siControl=60% ± 12, siCCDC15=61% ± 9, p=0.7254. **C.** Quantification of centrioles positive for Centrin-2 at the distal region of centrioles in control or CCDC15-depleted RPE1 cells based on (A). Error bars, SD. n>20 cells per experiment. Data represent mean value from two independent experiments per condition. siControl=80% ± 3, siCCDC15= 52% ± 0.85, p=0.0050. **D.** Representative U-ExM images of SFI1 protein in control, CCDC15 and POC5 siRNA-depleted RPE1 cells. Cells were expanded and immunostained with SFI1 (green) and tubulin (magenta) antibodies. Scale bar, 1 μm **E.** Quantification of centrioles positive for SFI1 at the distal region of centrioles in control and CCDC15-depleted RPE1 cells based on (D). Error bars, SD. n>20 cells per experiment. Data represent mean value from two independent experiments per condition. siControl=93.4%±0.1, siCCDC15=56.6%±2.2, p=0.0018; siControl=96.5%±1.5, siPOC5=96.4%±1.5, p=0.9625. **F.** Representative U-ExM images of coverages of CCDC15 in control, POC5, POC1B and FAM161A-depleted RPE1 cells. Cells were expanded and immunostained with CCDC15 (green) and tubulin (magenta) antibodies. Scale bar, 1 μm **G.** Quantification of CCDC15 coverage with respect to tubulin represented in (G). Error bars, SD. n>10 cells per experiment. Data represent mean value from three independent experiments per condition. siControl=1 ± 0.4; siPOC1B=0.44 ± 0.32; siPOC5=1.3 ± 0.53; siFAM161A=0.64 ± 0.24, p<0.0001. **H.** Representative U-ExM images of centrioles from RPE1 cells transfected with control siRNA or CCDC15 and POC1B siRNAs together. Cells were stained for CCDC15 and POC1B in green and tubulin in magenta. Scale bar, 1 μm **I.** Representative U-ExM images of defective centrioles in CCDC15 and POC1B co-depleted cells. Cells were co-stained with CCDC15 and POC1B in green and tubulin in magenta. Scale bar, 1 μm **J.** Centriole length quantification of (I). Error bars, SD. n>15 per experiment. Data represents mean value from two independent experiment. siControl: 1 ± 0.13, siCCDC15: 0.87 ± 0.11, siCCDC15/siPOC1B: 0.84 ± 0.10. **K.** Percentage of cells with defective centrioles for the indicated cells in CCDC15 and POC1B co-depleted cells. n>15 per experiment. Error bars, SD. Data represents mean value from two independent experiment. siControl=3.35% ± 1.1, siCCDC15=12% ± 0.28, siCCDC15/siPOC5=23.5% ± 1.13.

Nanoscale mapping of Centrin-2 revealed its dual localization at the central core region and the distal end of the centrioles (Le Guennec et al., 2020; Steib et al., 2020). Centrin-2 was recently described to form a complex with SFI1 to regulate centriole architecture and ciliogenesis (Laporte et al., 2022). Therefore, we examined whether CCDC15 depletion could affect the distal end localization of Centrin-2 and SFI1. Remarkably, analysis of the top and longitudinal views of centrioles showed that CCDC15 depletion resulted in a significant decrease in the percentage of cells with distal Centrin-2 and SFI-1 pools at the distal end of centrioles (Fig. 6A, 6C, 6D, 6E). To further confirm the specificity of this phenotype, we depleted the inner scaffold protein POC5 and examined its impact on centriolar localization of SFI1. As previously described, POC5 depletion did not alter percentage of cells with distal SFI1 centriolar pools (Fig. 6D, E) (Laporte et al., 2022). These results identify CCDC15 as a dual regulator of centriolar recruitment of inner scaffold protein POC1B and the distal end SFI1/Centrin complex.

Once we identified CCDC15 role in localizing inner scaffold proteins at the centriole, we next investigated the complementary effect, asking if inner scaffold proteins could regulate CCDC15 recruitment. To this end, we depleted the inner scaffold proteins POC5, POC1B and FAM161A using previously described siRNAs and validated their depletion by immunoblotting (Fig. S4F). Using U-ExM, we first quantified the CCDC15 centriolar coverage in these cells. While the centriolar coverage of CCDC15 increased upon POC5 depletion and decreased upon POC1B depletion, it remained unaltered upon FAM161A depletion (Fig. 6F, 6G). Consistently, the centrosomal abundance of CCDC15 decreased in POC1B or FAM161A-depleted cells and increased in POC5-depleted cells (Fig. S4G, S4H).

Since our results show that POC1B and CCDC15 rely on each other for their centriolar localization, we hypothesized that they might cooperate during regulation of centriole stability and length. To test this, we depleted CCDC15 by itself or together with POC1B and confirmed their efficient co-depletion by U-ExM analysis of cells stained for CCDC15 and POC1B (Fig. 6H). Although their co-depletion did not have an additive effect on centriole length shortening, the percentage of defective centrioles increased by about 2-fold in co-depleted cells relative to POC1B and CCDC15 depletion alone (Fig. 6H-K). Notably, the observed phenotypes in centriole wall breakage and shape were more severe in co-depleted cells (Fig. 6I). These results indicate that CCDC15/POC1B co-depletion enhances centriole architecture abnormalities.

### CCDC15 is required for assembly of signaling-competent primary cilium

Previous studies showed that the inner scaffold proteins including WDR90, Centrin-2 and POC5, as well as the distal SFI1/Centrin complex are required for proper cilium formation and function (Delaval et al., 2011; Hassan et al., 2019; Laporte et al., 2022; Prosser and Morrison, 2015; Steib et al., 2020). To investigate whether CCDC15 functions in cilium assembly, we depleted CCDC15 in RPE1 cells and quantified the percentage of ciliated cells upon 48-h serum starvation. While control cells ciliated at 70.15% ± 6.36%, CCDC15-depleted cells ciliated at 45.58% ± 0.81% (Fig. 7A, 7B). As quantified from immunofluorescence images and assessed by U-ExM, the cilia that formed in CCDC15-depleted cells were significantly shorter in length (Fig. 7A, 7C, 7D). Since CCDC15 was not efficiently depleted from the mother centriole, we investigated the link between CCDC15 absence and defective ciliogenesis by U-ExM, to select depleted cells. We co-stained CCDC15 with acetylated tubulin, which marks the cilium. While CCDC15-positive centrioles formed primary cilia, centrioles that lacked CCDC15 signal formed shorter cilia (Fig. 7D). This result further corroborates the role of CCDC15 during primary cilium assembly.

**Fig. 7.**
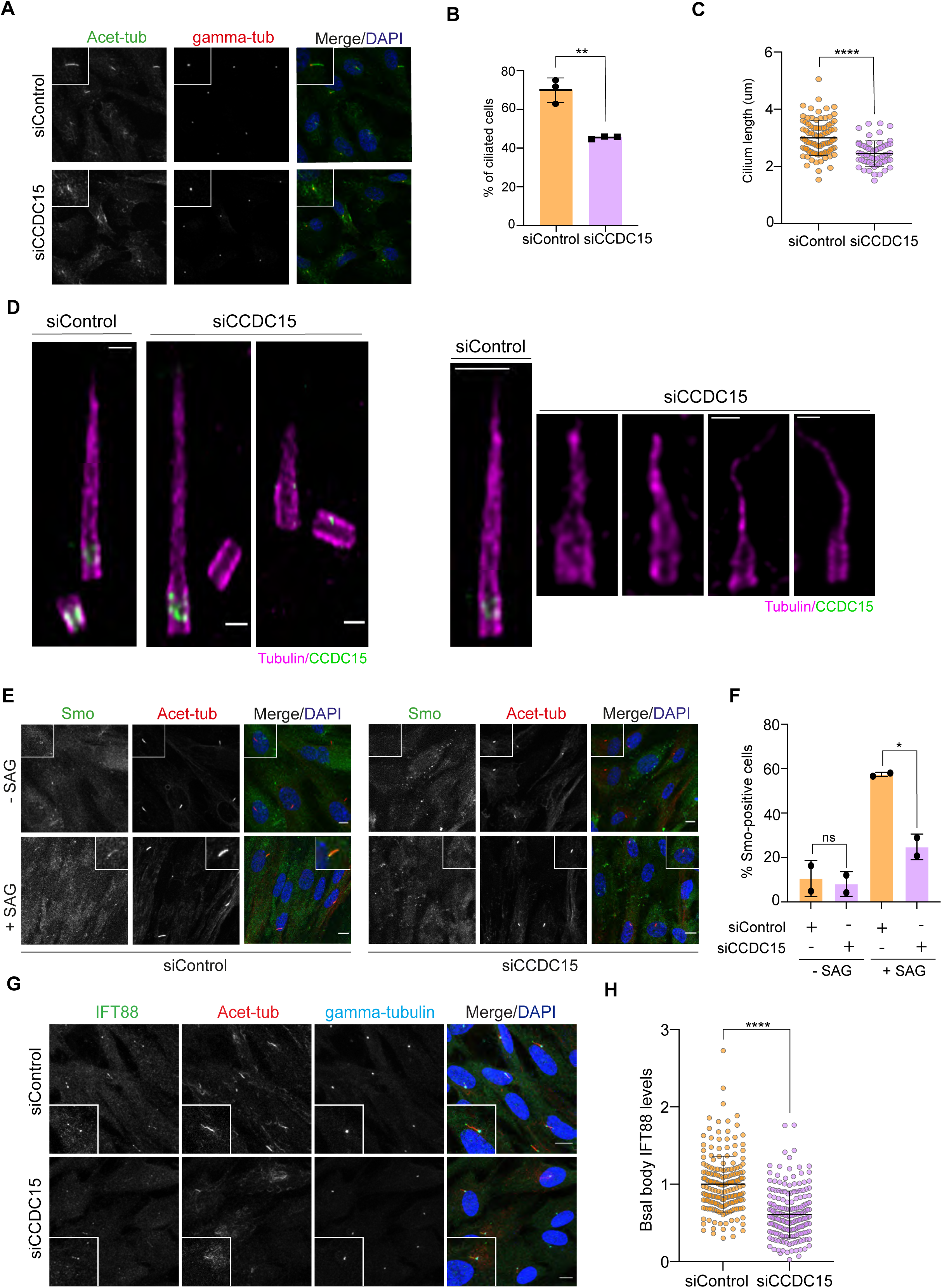
CCDC15 is required for assembly of signaling-competent primary cilium. **A.** Representative immunofluorescence images of ciliogenesis defects in RPE1 cells. Cells were transfected with control or CCDC15 siRNA. 72 h posttransfection, cells were serum starved for 24 h, fixed and immunostained for the primary cilium with acetylated tubulin antibody (Acet-tub) and the centrosome with gamma-tubulin antibody. DNA was visualized with DAPI. Scale bar, 10 μm. **B, C.** Quantification of ciliogenesis efficiency and length for (A). Error bars, SD. n>100 cells per experiment. Data represent mean value from three experiments per condition. Ciliation percentage: siControl=70% ± 6, siCCDC15=46% ± 0.8, p=0.0027. Cilium length: siControl=3 μm ± 0.7, siCCDC15=2.3 μm ± 0.4, p<0.0001. **D.** Representative U-ExM images of control and CCDC15-depleted RPE1 cells serum starved for 24 h. Centrioles and primary cilium were stained with tubulin (magenta) antibody and CCDC15 (green). Different ciliary defects associated with basal bodies efficiently depleted for CCDC15 were represented (panel on the right). Scale bar, 1 μm **E.** Representative immunofluorescence images of the effect of CCDC15 depletion on ciliary recruitment of Smo. Control and CCDC15 siRNA depleted RPE1 cells treated with 200 nM SAG for 24 h, fixed and stained for Smo, acetylated tubulin (Ac-tub), and DAPI. Scale bar, 10 μm **F.** Quantification of (E). Error bars, SD. Data represents mean value were the mean of two independent experiments. Smo-positive cells: siControl 57% ± 1, siCCDC15 25% ± 6, p=0.0173. **G.** Representative images of effect of CCDC15 depletion on basal body levels of IFT88. RPE1 cells were transfected with control or CCDC15 siRNA, fixed and stained for IFT88, acetylated tubulin and gamma-tubulin. DNA was visualized with DAPI. **H.** Quantification of (G). Error bars, SD. Data represents mean value were the mean of two independent experiments. p<0.0001

To assess the signaling competence of cilia that formed upon CCDC15 loss, we quantified cellular response to Hedgehog pathway activation. To this end, control and CCDC15-depleted cells were serum starved for 48 h to promote cilia formation and then stimulated for 24 h with 200 nM Smoothened agonist (SAG). We then characterized Smoothened (SMO) translocation efficacy to the cilia. As compared to control cells, CCDC15 depletion significantly decreased the percentage of SMO-positive cells (siControl 57.35 ± 0.997, siCCDC15 24.79 ± 5.798) (Fig. 7E, 7F). These results show that the cilia that forms upon CCDC15 depletion do not efficiently respond the Sonic hedgehog pathway.

Finally, we investigated whether the ciliary defects associated with CCDC15 loss were due to defects in centrioles’ ability to recruit key ciliogenesis factors. Given the important roles of the IFT-B machinery during cilium assembly and length regulation, we assessed how CCDC15 depletion affects basal body recruitment of the IFT-B component IFT88 in control and CCDC15-depleted cells (Mirvis et al., 2018; Nakayama and Katoh, 2018; Pazour et al., 2002). As compared to control cells, we observed a significant decrease in IFT88 levels at the basal body upon CCDC15 depletion, suggesting direct or indirect functions for CCDC15 in this process (Fig. 7G, 7H).

## Discussion

The inner scaffold of the centrioles recently emerged as an important centriolar sub-compartment that regulates centriole integrity, architecture, and function. Mutations affecting the inner scaffold were linked to retinal disorders (Langmann et al., 2010; Roosing et al., 2014; Weisz Hubshman et al., 2018; Ying et al., 2019). Despite its important functions and disease links, the full repertoire of inner scaffold proteins as well as how these proteins work together is not known. The *in vivo* proximity interactomes of Centrin-2 and POC5 identified previously characterized inner proteins and suggested new components, which therefore provides an important resource for future studies. Using U-ExM, coimmunoprecipitation and loss-of-function experiments, we identified CCDC15 as a novel centriole inner scaffold component critical for centriole length and integrity and, thereby, for its ability to template assembly of a functional primary cilium. CCDC15 stably associates with centrioles and interacts and co-localizes with known components of the centriole inner scaffold. Importantly, CCDC15 is a dual regulator of centriolar recruitment of the inner scaffold protein POC1B and the distal end SFI1/Centrin complex. Our findings suggest a model where CCDC15 functions as part of the inner scaffold to ensure assembly of full-length centrioles with proper architecture and function.

Loss-of-function experiments revealed CCDC15 functions during regulation of centriole size and integrity. Since inner scaffold proteins WDR90 and POC5 were also described to have similar functions, our results corroborate the role of the inner scaffold in these processes (Azimzadeh et al., 2009; Heydeck et al., 2020; Mercey et al., 2022; Steib et al., 2020). While WDR90 and POC5 depletion resulted in longer centrioles in interphase and shorter centrioles in prometaphase in U2OS cells, CCDC15 depletion resulted in shorter centrioles in RPE1 cells (Steib et al., 2020). Depletion of another centriole lumen protein HAUS6 also resulted in shorter centrioles in RPE1 cells (Schweizer et al., 2021). These results highlight the complexity of centriole length regulation as well as the necessity to investigate further in future studies how different inner scaffold proteins take part in this regulation.

To elucidate the molecular basis of CCDC15-associated centriolar defects, we assessed the extent of its interaction and localization with known inner scaffold proteins. We also examined whether they depend on each other for centriolar localization. Notably, CCDC15 and POC1B localizations to the inner scaffold were interdependent and their co-depletion exacerbated the centriolar defects compared to CCDC15 depletion alone. CCDC15 and POC1B might be involved in modulating each other’s turnover or interactions with other proteins, which needs to be investigated in future studies. POC1B was also reported to form a functional complex with CEP44 that ensures structural integrity of centrioles and their conversion to centrosomes (Atorino et al., 2020). Future studies that explore the relationship of CCDC15 to CEP44 and other central core proteins are required to uncover the precise mechanisms by which they operate at the central core of centrioles.

The distal end of the centrioles is directly engaged in ciliogenesis via removal of the CP110/CEP97 centriole cap complex and recruitment of ciliogenesis factors (Kim and Dynlacht, 2013; Sanchez and Dynlacht, 2016). For example, the inner scaffold protein Centrin-2 also localizes to the centriole distal end where it interacts with SFI1 and regulates centriole architecture as well as ciliogenesis by removing the CP110 cap (Laporte et al., 2022). Unlike Centrin-2, CCDC15 localization was limited to the central core of the centrioles, which is reminiscent of most of the known inner scaffold proteins (Hamel et al., 2017; Laporte et al., 2022; Le Guennec et al., 2020; Steib et al., 2020). Despite its central core localization, depletion of CCDC15 resulted in the loss of the distal pool of Centrin-2 without altering its localization to the central core. How CCDC15 regulates centriolar recruitment of distal end proteins without spatially occupying this region remains elusive. Perhaps CCDC15 transiently localizes to this region during centriole assembly, which might explain its presence in the proximity interactome of Centrin-2. Alternatively, CCDC15 might mediate the interaction between inner scaffold and distal end proteins via Centrin-2.

Our findings also revealed CCDC15 functions during assembly of the primary cilium with proper length. While defective ciliogenesis is a recurrent phenotype associated with depletion of proteins that are required for centriole structural integrity like WDR90, POC5 and POC1B, our understanding of the mechanisms by which these proteins regulate cilium assembly is still poorly understood (Mercey et al., 2022; Pearson et al., 2009; Schweizer et al., 2021; Steib et al., 2020). Since ciliogenesis requires elongation of microtubule doublets at the distal end of mother centrioles, it is tempting to speculate that centriolar defects associated with inner scaffold deregulation interferes with the ability of centrioles to template primary cilium assembly. Centriole architecture aberrations might lead to defective recruitment of ciliogenesis factors to the centrioles. In agreement, centriolar targeting of IFT88 was compromised in CCDC15-depleted cells. Moreover, SFI1/Centrin complex at the centriole distal end was shown to regulate ciliogenesis via removal of the centriole cap and organization of distal appendages (Laporte et al., 2022). Different stages of the ciliogenesis pathway needs to be studied in the future to fully uncover how inner scaffold regulates primary cilium assembly. In contrast to the primary cilium, the function of the inner scaffold has not yet been characterized during assembly and motility of motile cilia and flagellum. Since inner scaffold confers structural integrity to the centrioles when they are under mechanical stresses generated by ciliary motility, its future characterization in different cell types will provide important insight into mechanisms underlying its context-dependent functions.

Although centrioles in CCDC15-depleted cells were defective in cilium assembly, we did not observe any defect in canonical centriole duplication, which is initiated on the wall of the pre-existing centrioles at a single site during S phase. Alterations in centriole numbers were also not described in loss-of-function studies of other inner scaffold proteins such as WDR90 and POC5, suggesting that inner scaffold-regulated morphological and structural features do not play a role during canonical centriole duplication (Azimzadeh et al., 2009; Hamel et al., 2017). Intriguingly, CCDC15 depletion led to defects in centriole amplification induced by S phase arrest or PLK4 overexpression. It is possible that the aberrant centrioles associated with loss of the inner scaffold cannot withstand the forces exerted by the extra procentrioles during centriole amplification. It would therefore be interesting to study the role of the inner scaffold during centriole amplification in specialized cells such as multiciliated epithelia and olfactory cells.

Mutations in genes encoding the inner scaffold proteins POC5, POC1B, Centrin-2 and FAM161A were reported in human retinal disorders, which lead to photoreceptor degeneration and vision loss (Langmann et al., 2010; Roosing et al., 2014; Weisz Hubshman et al., 2018; Ying et al., 2019). Given its interactions with these proteins, CCDC15 might be a candidate gene for retinopathies. Future DNA sequencing of ciliopathy patients and *in vivo* studies are required to explore this possibility. Deregulation of centriole size has also been linked to diseases other than ciliopathies. For example, a screen for centrosome abnormalities in the NCI-60 panel of human cancer cell lines identified deregulation of centriole length as a recurrent feature of cancer (Marteil et al., 2018). Over-elongated centrioles were shown to enhance microtubule nucleation and chromosomal instability (Marteil et al., 2018). Interestingly, aberrant splicing of CCDC15 was implicated in Esophageal squamous cell carcinoma tumorigenesis (Tang et al., 2020). Therefore, uncovering the precise mechanisms by which centriole length is regulated by CCDC15 and its interactors might provide insight into tumorigenic mechanisms.

## Materials and Methods

### Plasmids

pDEST-GFP-CCDC15, pDEST-GFP-FAM161A, pDEST-FlagBirA*-CCDC15, pDEST-GFP-Centrin2, pLVPT2-V5-BirA*-SAS6, pLVPT2-V5-BirA*-POC5 and p pLVPT2-V5-BirA*-Centrin2 were generated by Gateway recombination between conor plasmids and the indicated Gateway destination plasmids. GFP-POC1B and mCherry-FAM161A were previously described in Le Guennec et al. 2020 (Le Guennec et al., 2020). Full-length CCDC15 was amplified by PCR and cloned into pCDH-EF1-mNeonGreen-T2A-Puro lentiviral expression plasmid.

### Cell culture, transfection and lentiviral transduction

Human telomerase immortalized retinal pigment epithelium cells (hTERT-RPE1, ATCC, CRL-4000) and inducible RPE1::Myc-PLK4 were cultured with Dulbecco’s modified Eagle’s Medium DMEM/F12 50/50 medium (Pan Biotech,Cat. # P04-41250) supplemented with 10% Fetal Bovine Serum (FBS, Life Technologies, Ref. # 10270-106, Lot # 42Q5283K) and1% penicillin-streptomycin (GIBCO, Cat. # 1540-122). Human embryonic kidney (HEK293T, ATCC, CRL-3216), and osteosarcoma epithelial (U2OS, ATCC, HTB-96) cells were cultured with DMEM medium (Pan Biotech, Cat. # P04-03590) supplemented with10% FBS and 1% penicillin-streptomycin. All cell lines were authenticated by Multiplex Cell Line Authentication (MCA) and were tested for mycoplasma by MycoAlert Mycoplasma Detection Kit (Lonza). U2OS cells were transfected with the plasmids using Lipofectamine 2000 and according to the manufacturer’s instructions (Thermo Fisher Scientific). For pulldown experiments, total of 10 µg of plasmids were transfected to HEK293T cells using 1mg/ml polyethylenimine, MW 25 kDa (PEI). For microtubule depolymerization experiments, cells were treated with 10 µg/ml nocodazole (Sigma-Aldrich, Cat. #M1404) or vehicle (dimethyl sulfoxide) for one hour at 37°C. For cell synchronization experiments, 5 µM (+)-S-trityl-L-cysteine (STLC) (Alfa-Aesar, Cat. #2799-07-7) was used for 16 h at 37°C. For hydroxyurea treatment, U2OS cells are treated with 4 mM hydroxyurea for 48 hours. Lentivirus were generated using pLVPT2-V5-BirA*-POC5 and p pLVPT2-V5-BirA*-Centrin2 plasmids as transfer vectors. HEK293T cells were transduced with the indicated lentivirus and selected with 2 µg/ml puromycin for four to six days until all the control cells died.

### siRNA transfections

Cells were seeded onto coverslips at 30-40% confluency and transfected with 50 nM of siRNA in two sequential transfections using Lipofectamine RNAiMAX (Life Technologies) in OPTI-MEM (Life Technologies) according to the manufacturer’s instructions. Depletion of proteins was confirmed at 48 h, 72 h or 96 h after transfection by immunofluorescence and immunoblotting. CCDC15 was depleted using an siRNA with sequence 5′-GCAGUACCUGAGACAUAGAtt-3′ (Ambion, Cat. # s36888). For depletion of POC5, POC1B and FAM161A, published siRNA sequences were used (Atorino et al., 2020; Di Gioia et al., 2012; Steib et al., 2020).

### Immunofluorescence, antibodies, and microscopy

Cells were grown on coverslips, washed twice with PBS and fixed with either ice cold methanol at -20°C for 10 minutes or 4% PFA in cytoskeletal buffer (10 mM PIPES, 3 mM MgCl_2_, 100 mM NaCl, 300 mM sucrose, pH 6.9) supplemented with 5 mM EGTA and 0.1% Triton X for 15 minutes at 37°C. After washing twice with PBS, cells were blocked with 3% BSA in PBS + 0.1% Triton X-100 and incubated with primary antibodies in blocking solution for 1 hour at room temperature. Cells were washed three times with PBS and incubated with secondary antibodies and DAPI (Thermo Scientific) at 1:2000 for 45 minutes at room temperature. Following three washes with PBS, cells were mounted using Mowiol mounting medium containing N-propyl gallate (Sigma-Aldrich). Primary antibodies used for immunofluorescence were rabbit anti CCDC15 (1:1000, Invitrogen PA5-59184), mouse anti Centrin-2 (1:1000, Sigma Aldrich 04-1624), mouse anti *γ*-tubulin (GTU88, 1:5000, Sigma Aldrich, T6557), mouse anti Cep164 (1:1000, Santa Cruz Biotechnology, sc515403), mouse anti SAS6 (1:1000, Santa Cruz Biotechnology, sc81431), mouse anti rootletin (1:500, Santa Cruz Biotechnology, sc374056), mouse anti glutamylated-tubulin (GT335, 1:1000, Adipogen, AG-20B-0020), mouse anti Centrin-3 (1:1000, Abnova, H8141-3EG), mouse anti V5 (1:1000, Invitrogen, 46-0705), mouse anti GFP (1:1000, Life Technologies, A-11120), rabbit anti CEP63 (1:1000, Millipore, 06-1292), rabbit anti CEP152 (1:1000, Bethyl, A302-479), mouse anti *α*-tubulin (DM1A, 1:10000, Sigma Aldrich, T926-2ML), rabbit anti POC5 (1:750, Invitrogen, A303-341A), rabbit anti POC1B (1:1000, Thermo Fisher Scientific, PA5-24495), rabbit anti FAM161(1:1000, Invitrogen, PA5-56935), rabbit anti Centrin1 (1:1000, Proteintech, 127941AP), rabbit anti PCM1 (1:1000, homemade), rabbit anti-CP110 (Proteintech-12780-1-AP) and mouse anti acetylated tubulin (1:10000, Sigma Aldrich, 059M4812V). Secondary antibodies used for immunofluorescence experiments were AlexaFluor 488-, 568- or 633-coupled (Life Technologies) and they were used at 1:2000. Biotinylated proteins were detected with streptavidin coupled to Alexa Fluor 594 (Thermo Fisher).

Quantitative immunofluorescence of centrosomal and ciliary levels of proteins was performed by acquiring a z-stack of cells using identical gain and exposure settings, determined by adjusting settings based on the fluorescence signal in the control cells. The z-stacks were used to assemble maximum-intensity projections. The centrosome regions in these images were defined by centrosomal marker gamma-tubulin or centrin staining for each cell, and the total pixel intensity of a circular 3 μm^2^ area centered on the centrosome in each cell was measured using ImageJ and defined as the centrosomal intensity. The ciliary regions in these images were defined by ciliary markers ARL13B or acetylated tubulin staining for each cell. Total pixel intensity of fluorescence within the region of interest was measured using ImageJ (National Institutes of Health, Bethesda, MD) (Schneider et al., 2012). Background subtraction was performed by quantifying fluorescence intensity of a region of equal dimensions in the area proximate to the centrosome or cilium. Statistical analysis was done by normalizing these values to their mean. Efficiency of primary cilium formation was quantified by counting the total number of cells, and the number of cells with primary cilia, as detected by DAPI and acetylated tubulin staining, respectively. The cilium length was quantified using acetylated tubulin as the primary cilium marker.

### Fluorescence recovery after photobleaching

FRAP experiments were performed with Leica SP8 confocal microscope using FRAP module. U2OS cells were incubated with 10% FBS in DMEM and RPE1 cells were incubated with 10% FBS in DMEM-12 and kept at 37 °C with 5% CO2. ROI was set to include either one centriole or two centrioles. A z-stack of 8 µm with 01 µm step size was taken during pre and post bleaching for both one and two centriole FRAP experiments. Bleaching was done 4 iterations with 488 Argon laser with 100% power. Maximal projection of the files was performed in Leica LAS X software and analysis was done in ImageJ. Recovery graph quantifications, t-half and mobile pool quantifications were done using the equations as described (Sprague et al., 2004).

### Immunoprecipitation

HEK293T cells were co-transfected with indicated plasmids. 48 h post-transfection, cells were washed and lysed either with LAP200 buffer (50 mM HEPES, pH 7.4, 100 mM KCl, 1 mM EGTA, 1 mM MgCl2, 10% glycerol, 0.3% NP40 freshly supplemented with protease inhibitors, 10 µg/ml Leupeptin, Pepstatin and Chymostatin, 1 mM PMSF) or Flag pulldown buffer (50 mM HEPES pH 8, 100 mM KCl, 2 mM EDTA, 10% glycerol, 0.1 % NP-40 freshly supplemented with 1 mM DTT and protease inhibitors, 10 µg/ml Leupeptin, Pepstatin and Chymostatin, 1 mM PMSF) for GBP and Flag beads, respectively for 30 min. Lysates were centrifuged at 13000 rpm for 10 min at +4°C and supernatants were transferred to a tube. 100 μl from each sample was saved as input. The rest of the supernatant was immunoprecipitated with GBP beads (homemade) or Anti-FLAG M2 agarose beads (Sigma Aldrich) overnight at +4°C. After washing 3x with indicated lysis buffer, samples were resuspended in SDS containing sample buffer. The samples are resolved in SDS-PAGE gels and transferred onto nitrocellulose membranes, blocked with TBST in 5% milk for 1 hour at room temperature. Blots were incubated with primary antibodies diluted in 5% BSA in TBST overnight at 4 °C and washed with TBST containing 1% Tween-20 three times for 10 minutes. They are incubated with secondary antibodies at room temperature for 1 hours and washed three times with TBST containing 1% Tween-20. The blots were visualized with the LI-COR Odyssey® Infrared Imaging System and software at 169 µm (LI-COR Bioscience). The primary antibodies used for immunoblotting are rabbit anti GFP (1:500, homemade), Mouse anti V5 (1:500, Invitrogen, 46-0705), rabbit anti Flag (1:5000, Proteintech, 20543-1-AP) and mouse anti Flag (1:5000, Proteintech, 66008-3-Ig). Secondary antibodies used for western blotting experiments were IRDye680- and IRDye 800-coupled and were used at 1:10000 (LI-COR Biosciences).

### Cell Cycle Analysis

RPE1 cells were seeded to 12 well plates and treated with either control siRNA or CCDC15 siRNA for 96 h. Cells were collected with trypsin, centrifuged at 300 g for 5 minutes and washed with 1X PBS. 50 μl of PBS was left at the bottom of the tube and the pellet is resuspended in the PBS. The resuspended cells were added onto the 1 mL of 70% ethanol drop by drop while gently vortexing. Fixed cells were incubated at -20°C for 3 hours prior to staining. Cells were centrifuged at 300g for 5 minutes, washed with 1X PBS and stained with 200 μl Muse^TM^ Cell Cycle Reagent (Luminex Corporation) for 30 minutes before analysis. Samples are run in The Guava® Muse® Cell Analyzer.

### UltraStructure Expansion Microscopy (U-ExM) and image analysis

U-ExM was performed as previously described (Gambarotto et al., 2021). Briefly, RPE1 cells were grown on 12 mm coverslips in 24 well plate. Coverslips were incubated in 1.4% formaldehyde / 2% acrylamide (2X FA / AA) solution in 1X PBS for 5 h at 37°C. Cells are embedded into a gel prepared with Monomer Solution (for one gel, 25 μl of sodium acrylamide, stock solution at 38% (w/w) diluted with nuclease-free water, 12.5 μl of AA, 2.5 μl of BIS and 5 μl of 10X PBS) supplemented with TEMED and APS (final concentration of 0.5%) for one hour at 37°C. Denaturation was performed at 95°C for 90 minutes and gels were stained with primary antibodies for 3 hours at 37°C. Gels were washed three times 10 min at RT with 1X PBS with 0.1% Triton-X (PBS-T). Secondary antibody incubation is carried out for 2h30 at 37°C and gels washed with three times 10 min washes in PBS-T at RT. Gels were expanded in 100 ml dH_2_O three times before imaging. The diameter of the gels is measured with a ruler and the expansion factor is calculated by dividing the diameter to 12mm. Gels were cropped into pieces, and they are mounted to 24mm coverslips coated with Poly-D-lysine. The images are taken with Leica SP8 confocal microscope with 0.30 μm z-intervals and deconvolved in Huygens software. The primary antibodies used in these experiments are rabbit anti CCDC15 (1:500, Invitrogen, PA5-59184), rabbit anti-POC5 (1:500, Invitrogen, A303-341A), rabbit anti-POC1B (1:500, Thermo Fisher Scientific, PA5-24495), rabbit anti-FAM161A (1:500, Invitrogen, PA5-56935), rabbit anti-SFI1 (1:500, Proteintech, 13550-1-AP), mouse anti-Centrin-2 (1:500, Sigma Aldrich, 04-1624), rabbit anti-CEP164 (1:500, Proteintech, 22227-1-AP), AA344 (1:250, scFv-S11B, Beta-tubulin) and AA345 (1:250, scFv-F2C, Alpha-tubulin). The secondary antibodies used in these experiments are goat anti-rabbit 488 and goat anti mouse H+L 568 (Life Technologies) at 1:1000 and goat anti guinea pig 568 (1:1000, Invitrogen, A-11075). For U-ExM data, the RPE1 cells that are in G1 phase are quantified. Coverages of the proteins in U-ExM images are calculated as previously published in Le Guennec et al., 2020 (Le Guennec et al., 2020). For diameter measurements, a straight line from the exterior to the interior of the centriole was drew displaying a resolved signal for both tubulin and the core protein. The position value of the core protein’s maximum fluorescence intensity was aligned to the position of the corresponding tubulin maximal intensity for tubulin measurement. The values for the distance were plotted and analysed in GraphPadPrism7.

For measurement of centriole diameter at distinct positions, a straight line spanning the centriole is drawn within the distal, middle and proximal regions with respect to the positions of POC5, POC1B and FAM161A. The distal region is taken as the portion of the centriole above the staining of POC5, FAM161A or POC1B and the proximal region above them. The middle part is defined as the middle of each centriole. The plot profile toll of Fiji is used to gather the data and it was analysed by the script described in Le Guennec et al., 2020 (Le Guennec et al., 2020).

### Biotin-streptavidin affinity purification

For the BioID experiments, HEK293T stably expressing V5-BirA*-POC5 or V5-BirA*-Centrin-2 were grown in 5×15 cm plates were grown in complete medium supplemented with 50 μM biotin for 18 h. Following biotin treatment, cells were lysed in lysis buffer (20 mM HEPES, pH 7.8, 5 mM K-acetate, 0.5 mM MgCI2, 0.5 mM DTT, protease inhibitors) and sonicated. An equal volume of 4°C 50 mM Tris (pH 7.4) was added to the extracts and insoluble material was pelleted. Soluble materials from whole cell lysates were incubated with Streptavidin agarose beads (Thermo Scientific). Beads were collected and washed twice in wash buffer 1 (2% SDS in dH2O), once with wash buffer 2 (0.2% deoxycholate, 1% Triton X-100, 500 mM NaCI, 1 mM EDTA, and 50 mM Hepes, pH 7.5), once with wash buffer 3 (250 mM LiCI, 0.5% NP-40, 0.5% deoxycholate, 1% Triton X-100, 500 mM NaCI, 1 mM EDTA and 10 mM Tris, pH 8.1) and twice with wash buffer 4 (50 mM Tris, pH 7.4, and 50 mM NaCI). 10% of the sample was reserved for Western blot analysis and 90% of the sample to be analyzed by mass spectrometry was washed twice in 50 mM NH_4_HCO_3_.

### Biotin-streptavidin affinity purification and mass spectrometry

HEK293T cells stably expressing V5BirA*-POC5 or V5BirA*-Centrin-2 were generated by lentiviral transduction. For mass spectrometry analysis, each cell type was grown in 5×15 cm plates in DMEM medium supplied with 10% FBS and 1% penicillin-streptomycin and with 50 μM biotin for 18 h. Following biotin treatment, cells were lysed in lysis buffer (20 mM HEPES, pH 7.8, 5 mM K-acetate, 0.5 mM MgCI2, 0.5 mM DTT, protease inhibitors) and sonicated. An equal volume of 4°C 50 mM Tris (pH 7.4) was added to the extracts and insoluble material was pelleted. Soluble materials from whole cell lysates were incubated with Streptavidin agarose beads (Thermo Scientific). Beads were collected and washed twice in wash buffer 1 (2% SDS in dH2O), once with wash buffer 2 (0.2% deoxycholate, 1% Triton X-100, 500 mM NaCI, 1 mM EDTA, and 50 mM Hepes, pH 7.5), once with wash buffer 3 (250 mM LiCI, 0.5% NP-40, 0.5% deoxycholate, 1% Triton X-100, 500 mM NaCI, 1 mM EDTA and 10 mM Tris, pH 8.1) and twice with wash buffer 4 (50 mM Tris, pH 7.4, and 50 mM NaCI). 10% of the sample was reserved for Western blot analysis and 90% of the sample to be analyzed by mass spectrometry was washed twice in 50 mM NH_4_HCO_3_. Mass spectrometry analysis were performed as previously described (Arslanhan et al., 2021; Gurkaslar et al., 2020).

### Mass Spectrometry Data Analysis

For identification of high confidence proximity interactomes for Centrin-2 and POC5, data from three biological replicates for V5-BirA*-Centrin-2, four biological replicates for V5-BirA*-POC5 were used along with control V5BirA* data as published in Arslanhan et al. 2021 (Arslanhan et al., 2021). For mass spectrometry analysis, Normalized Spectral Abundance Factor (NSAF) values were generated for each protein by dividing each Peptide Spectrum Match (PSM) value by the total PSM count in that dataset. The fold change was calculated by dividing the NSAF values of POC5 and Centrin-2 interactors to their NSAF values in control datasets. The NSAF values equal to 0 in the control condition was replaced with the smallest NSAF values represented in the dataset. Proteins with log^2^ NSAF value greater than 1 as well as proteins identified in at least two biological replicates were accounted.

Furthermore, the remaining proteins were submitted to CRAPome (https://reprint-apms.org), which is a contaminant repository for mass spectrometry data collected form affinity purification experiments and a list with contaminancy percentage (%) was calculated (Mellacheruvu et al., 2013). Proteins with contaminancy percentage more than 30% were considered as a contaminant and removed. The interaction maps were drawn with 480 and 68 interactors for Centrin-2 and POC5, respectively. The GO terms for the proximity interactors were determined by using Database for Annotation, Visualization, and Integrated Discovery (DAVID).

For Fig. S1E and S1F, we performed the network analyses for all the proximity interactors of Centrin-2 and POC5, and high-confidence interactors ranked by their relative fold change in V5-BirA*-Centrin-2 and V5-BirA*-POC5 dataset versus V5-BirA* dataset. The interaction networks of these proteins are plotted using STRING database, and the map is visualized by CytoScape. The functional clusters and GO categories for these clusters are determined with the Clustering with Overlapping Neighborhood Expansion (ClusterONE) plug-in of Cytoscape and BinGO plug-ins (P < 0.05). GO terms were determined by using Database for Annotation, Visualization, and Integrated Discovery (DAVID). The network output file was visualized using Cytoscape 3.7.2 (Nepusz et al, 2012).

### Statistical analysis

Statistical results, average and standard deviation values were computed and plotted by using Prism (GraphPad, La Jolla, CA). Student’s t-test was applied to compare the statistical significance of the measurements. Following key is followed for asterisk placeholders for *p*-values in the figures: \**P* < 0.05, \*\**P* < 0.01, \*\*\**P* < 0.001, **** *P* < 0.0001

## Acknowledgements

We acknowledge the Firat-Karalar lab members for insightful discussions regarding this work and Koç University Proteomics Facility (KUPAM), Büşra Akarlar and Nurhan Özlü for their help for mass spectrometry experiments. This project has received funding from the European Research Council Starting grant agreement No 679140 to E.N.F, EMBO Installation Grant and Young Investigator Award to E.N.F, TUBITAK BIDEB 120C148 grant to E.N.F and the Swiss National Science Foundation (SNSF) PP00P3_157517 and 310030_205087 to P.G and V.H.

---

**Fig. S1.**
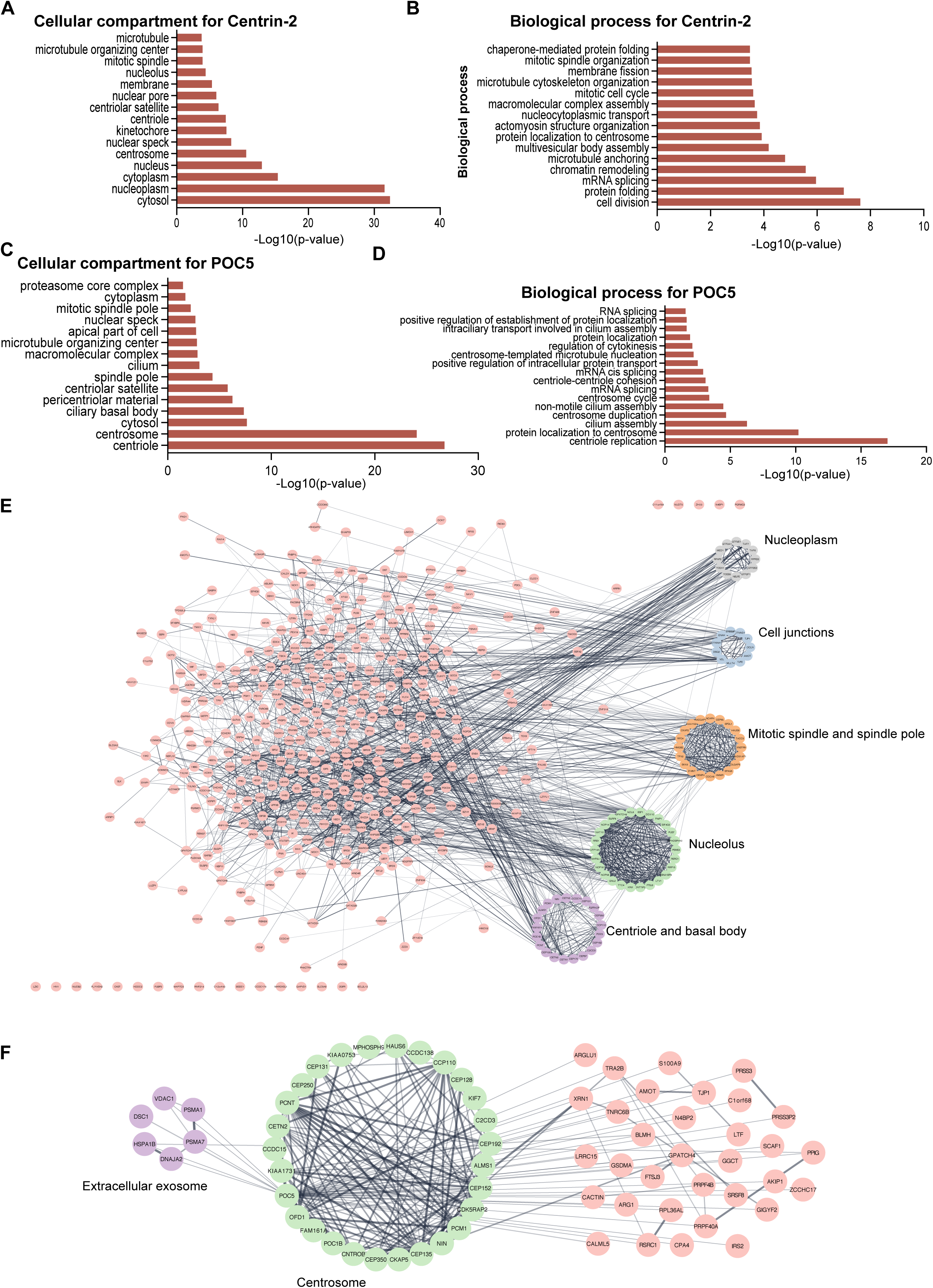
Gene enrichment analysis of the POC5 and Centrin-2 proximity interactomes. **A, B.** GO-enrichment analysis of Centrin-2 proximity interactors based on their cellular compartments and biological process. The x-axis represents the log-transformed p-value (Fisher’s exact test) of GO terms. **C, D.** GO-enrichment analysis of POC5 proximity interactors based on their cellular compartments and biological process. The x-axis represents the log-transformed p-value (Fisher’s exact test) of GO terms. **E, F.** Centrin-2 and POC5 proximity interactome maps. High confidence proximity interactors of POC5 and Centrin-2 were determined by using NSAF and CRAPome banalysis. The interaction map was generated using STRING protein interaction database and the proximity interactome of Centrin-2 were drawn in CytoScape. The clusters were determined by the ClusterONE plug-in CytoScape.

**Fig. S2.**
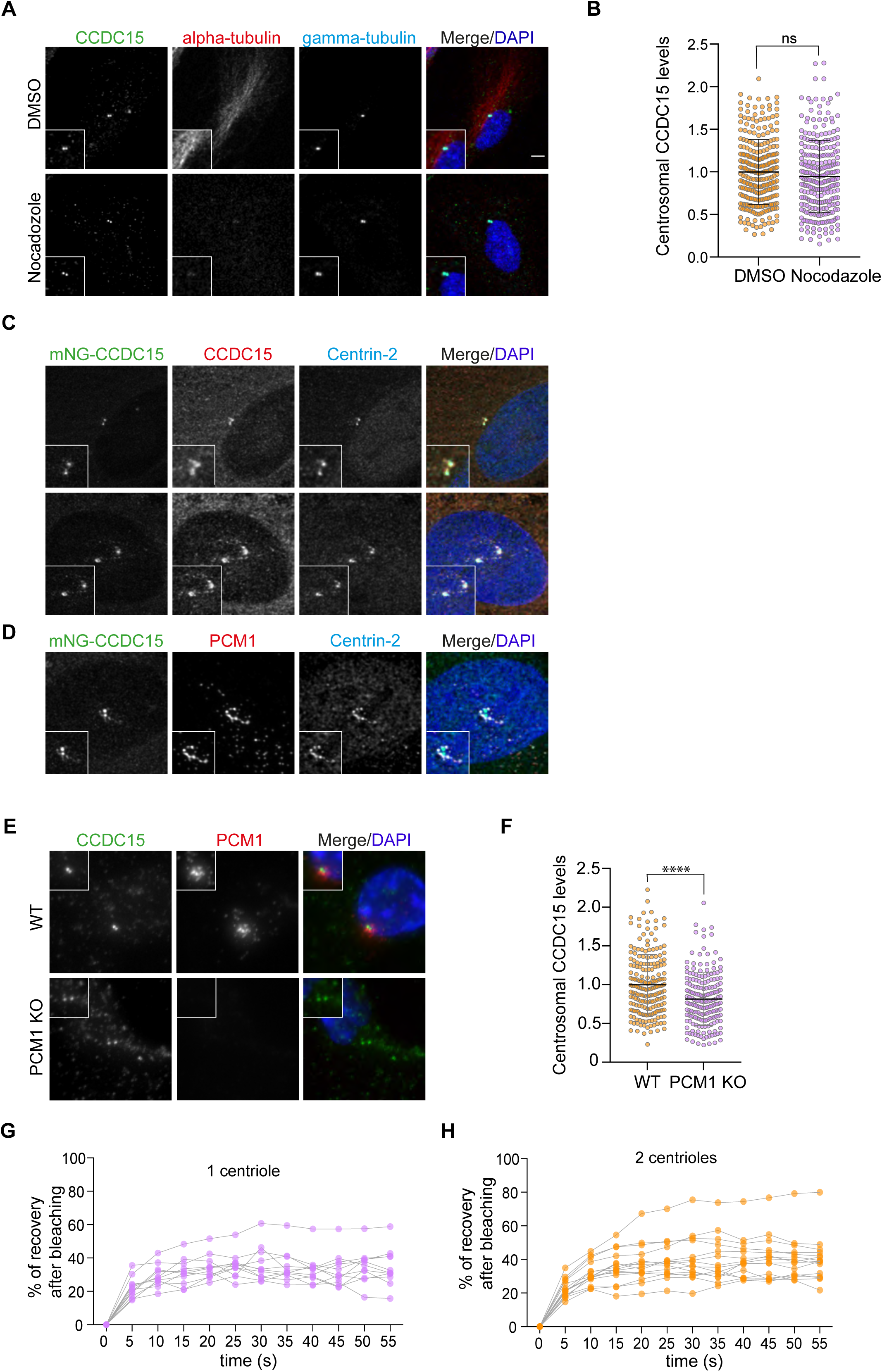
Analysis of CCDC15 localization. **A.** Centriolar recruitment of CCDC15 does not depend on microtubules. RPE1 cells were treated with DMSO (vehicle control) or nocodazole. Cells were fixed and stained for CCDC15, alpha-tubulin and gamma-tubulin. DNA was visualized with DAPI. **B.** Quantification of (A). Error bars, SD. n>100 cells per experiment. Data represent mean value from two independent experiments per condition. siControl=1±0.4, siCCDC15=0.95±0.43, p=0.1113. **C.** Localization of CCDC15 in transiently transfected RPE1 cells. U2OS cells were transfected with mNeonGreen (mNG)-CCDC15, fixed and stained for CCDC15 and Centrin-2. DNA was visualized with DAPI. **D.** Localization of CCDC15 in transiently transfected U2OS cells. U2OS cells were transfected with mNeonGreen (mNG)-CCDC15, fixed and stained for CCDC15 and Centrin-2. DNA was visualized with DAPI. **E.** Role of centriolar satellites in centrosomal targeting of CCDC15. RPE1 wild-type (WT) and satellite-less PCM1 KO cells were fixed and stained for CCDC15, PCM1 and DNA. **F.** Quantification of (E). n>100 cells per experiment. Data represent mean value from two independent experiments per condition. RPE1 WT=1±0.4, RPE1 PCM1 KO=0.82±0.34. p<0.0001 **G, H.** Percentage of recovery graphs of individual FRAP experiments in 1 centriole (G) and 2 centrioles (H). n=11 for 1 centriole, n=15 for 2 centrioles.

**Fig. S3.**
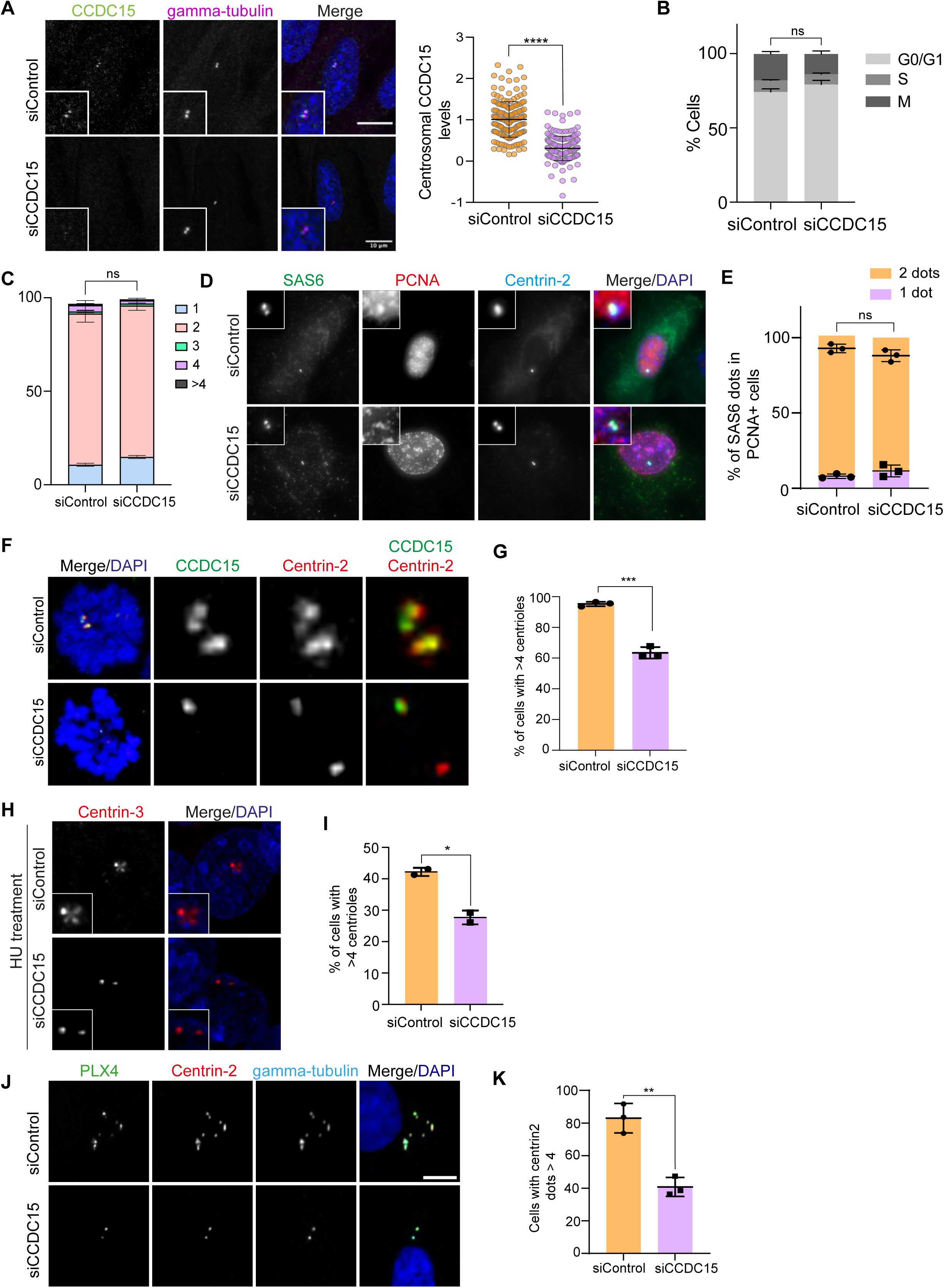
CCDC15 is required for centriole amplification, but not canonical centriole duplication. **A.** Immunofluorescence validation of CCDC15 depletion by siRNA treatment in RPE1 cells. RPE1 cells were transfected with control and CCDC15 siRNAs. 96 h after transfection, cells were fixed with methanol and stained for CCDC15 and gamma-tubulin. Error bars, SD. n>100 cells per experiment. Data represent mean value from two experiments per condition. siControl:1± 0.43, siCCDC15:0.31± 0.29 **B.** Cell cycle profile of control and CCD15-depelted RPE1 cells. RPE1 cells were transfected with control and CCDC15 siRNAs. 96 h after transfection, cellswere fixed with ethanol and stained with Muse Cell Cycle kit. Error bars, SD. Data represents mean value from two independent experiments per condition. **C.** Quantification of centriole number in control or CCDC15 siRNA-transfected asynchronous RPE1 cells. Error bars, SD. n>100 cells per experiment. Data represents mean value from two independent experiments per condition. **D.** Representative immunofluorescence images of control and CCDC15-depleted cells stained for SAS6, PCNA and Centrin-2. DNA was visualized with DAPI. **E.** Quantification of SAS6 dots in PCNA-positive cells in (D). Error bars, SD. n>50 cells per experiment. Data represent mean value from three experiments per condition. SAS6 2 dots: siControl=93%±3, siCCDC15=88%±4, p=0.1565; SAS6-1 dot: siControl=8%±1, siCCDC15= 12%±4, p=0.2172. **F.** Representative images of centrioles in control and CCDC15-depleted RPE1 cells synchronized by STLC treatment. Cells were transfected with control and CCDC15 siRNA and treated with 50 µM STLC for 18 h before fixation. Cells were then stained for CCDC15 and Centrin-2. The DNA was visualized with DAPI. **G.** Quantification of cells with more than 4 centrioles based on (F). Error bars, SD. n>100 cells per experiment. Data represent mean value from three experiments per condition. siControl=95%±1, siCCDC15=63%±4, p=0.0002. **H.** CCDC15 depletion compromises S phase-arrest overduplication of centrioles. U2OS cells were transfected with control siRNA or CCDC15 siRNA and arrested in S phase by hydroxyurea treatment for 48 h. Cells were than stained with CCDC15 and Centrin-3. DNA was visualized with DAPI. **I.** Quantification of cells with more than 4 centrioles based on (H). Error bars, SD. n>100 cells per experiment. Data represent mean value from two experiments per condition. siControl=42%±1, siCCDC15=28%±2, p=0.0148. **J.** CCDC15 depletion compromises PLK4-induced centriole amplification. RPE-1 cells stably expressing Tet-inducible Plk4 were depleted of CCDC15 by siRNA for 72 h then treated with doxycycline for 18 h to induce Plk4 expression. Cells were fixed and stained for PLK4, Centrin-2 and gamma-tubulin. DNA was visualized with DAPI. **K.** Quantification of cells with more than 4 centrioles dots based on (J). Error bars, SD. n>100 cells per experiment. Data represent mean value from three experiments per condition. siControl=83%±9, siCCDC15=41%±6, p=0.0024.

**Fig. S4.**
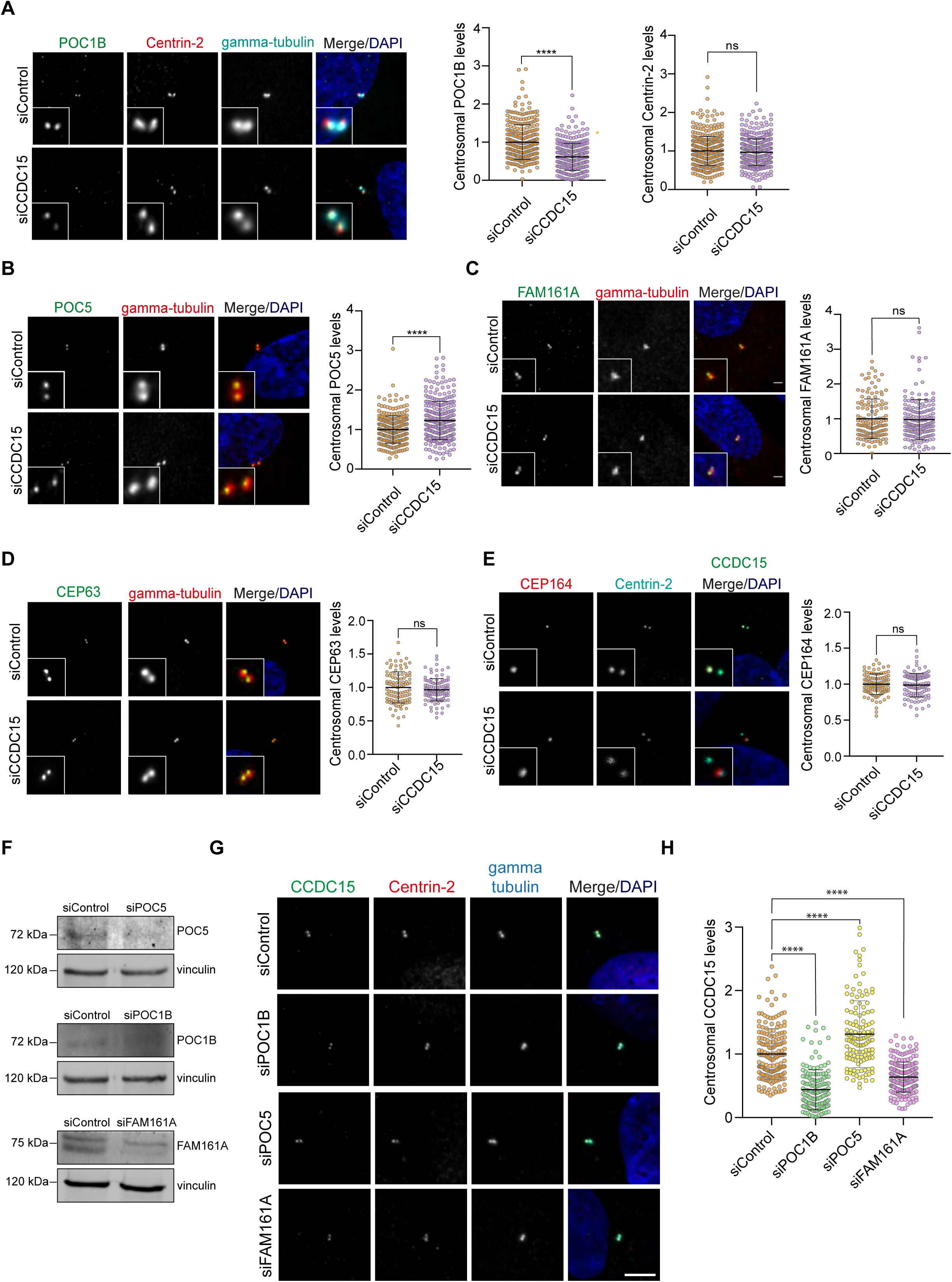
CCDC15 and inner scaffold proteins depend on each other for centrosomal abundance. **A-E.** Representative images and quantification of effect of CCDC15 depletion on centrosomal levels of (A) POC1B and Centrin-2, (B) POC5, (C) FAM161A, (D) CEP63 and (E) CEP164. RPE1 cells were fixed 96 h posttransfection with control or CCDC15 siRNA and stained for the indicated proteins. DNA was visualized with DAPI. Error bars, SD. n>100 cells per experiment. Data represent mean value from three experiments per condition. **F.** Validation of siRNA-mediated depletion of POC5, POC1B and FAM161A in RPE1 cells. Cells were transfected with control or POC5, POC1B and FAM161A siRNAs and extracts from these cells were immunoblotted for the indicated proteins and vinculin as loading control. **G.** Representative images of effect of POC5, POC1B or FAM161A depletion on centrosomal levels of CCDC15. RPE1 cells were fixed 96 h posttransfection with the indicated siRNAs and stained for CCDC15, Cenrin-2 and gamma-tubulin. DNA was visualized with DAPI. **H.** Quantification of CCDC15 centrosomal intensity based on (G). Error bars, SD. n>100 cells per experiment. Data represent mean value from three experiments per condition.

**Table S1. Centrin-2 and POC5 proximity interactor analysis**

Tab 1. Raw data and NSAF values of POC5, Centrin-2 and control AP/MS experiments. Columns indicate the spectral counts of each proteins.

Tab 2. POC5 proximity interactors identified in at least two biological replicates with spectral counts greater than one

Tab 3. POC5 proximity interactors after CRAPome contaminant removal

Tab 4. List of 68 proteins identified as high confidence interactors of POC5 with log^2^ NSAF value greater than 1

Tab 5. Centrin-2 proximity interactors identified in at least two biological replicates with spectral counts greater than one

Tab 6. Centrin-2 proximity interactors after CRAPome contaminant removal

Tab 7. List of 480 proteins identified as high confidence interactors of POC5 with log^2^ NSAF value greater than 1

**Table S2. GO enrichment analysis of high confidence interactors for POC5 and Centrin-2**

Tab 1. GO categories for cellular compartment and biological processes for Centrin-2 with p-values and -Log10 (p-value) that correspond to Fig. S2A and S2B.

Tab 2. GO categories for cellular compartment and biological processes for POC5 with p-values and -Log10 (p-value) that correspond to Fig. S2C and S2D.

**Movie S1. CCDC66 dynamic localization during cell cycle**

RPE1 cells stably expressing mNeonGreen-CCDC15 were imaged with confocal microscopy every 2 minutes. Scale bar: 1 µm

